# Aging-associated long non-coding RNA boosts longevity and reduces the ribosome content of non-dividing fission yeast cells

**DOI:** 10.1101/2024.03.25.586524

**Authors:** Shajahan Anver, Ahmed Faisal Sumit, Xi-Ming Sun, Abubakar Hatimy, Konstantinos Thalassinos, Samuel Marguerat, Nazif Alic, Jürg Bähler

## Abstract

Genomes produce widespread long non-coding RNAs (lncRNAs) of largely unknown functions. We characterize *aal1* (aging-associated lncRNA) which is induced in quiescent cells of fission yeast. Deletion of *aal1* shortens the chronological lifespan of non-dividing cells, while ectopic overexpression of *aal1* prolongs their lifespan, indicating that this lncRNA acts *in trans*. The overexpression of *aal1* leads to the repression of ribosomal protein genes and inhibition of cell growth, and *aal1* genetically interacts with coding genes functioning in protein translation. The *aal1* RNA localizes to the cytoplasm and associates with ribosomes. Notably, *aal1* deletion or overexpression is sufficient to increase or decrease the cellular ribosome content. The *rpl1901* mRNA, encoding a ribosomal protein, is a binding target of *aal1.* The levels of *rpl1901* are reduced ∼2-fold by *aal1,* which is critical and sufficient to extend the lifespan. Remarkably, the expression of *aal1* lncRNA in *Drosophila* triggers an extension of fly lifespan. We propose that *aal1* reduces the ribosome content by decreasing the levels of Rpl1901, thus attenuating protein translation and promoting longevity. Although the *aal1* lncRNA itself is not conserved, its effect in flies raises the possibility that animals feature related mechanisms that modulate aging, based on the conserved translational machinery.

## Introduction

Aging is a complex, multifactorial process leading to a gradual decline in biological function over time^1^. Old age is a shared root cause for complex diseases such as cancer, neurodegeneration, and cardiovascular or metabolic disorders^2^. Aging is highly plastic: simple genetic, nutritional, or pharmacological interventions in model organisms can extend lifespan^3^. But a major challenge for aging research remains to understand all factors affecting lifespan. Aging-related processes are remarkably conserved from yeast to humans, with simple, short-lived model organisms remaining the main platform to discover and dissect factors that modulate aging processes^4^.

Using fission yeast (*Schizosaccharomyces pombe*) as a simple model for cellular aging, we and others have analyzed genetic and environmental factors determining chronological lifespan (CLS)^5–8^. CLS is a complex trait defined as the time cells survive under limiting nutrients in a non-dividing state, which enlightens the aging of post-mitotic and quiescent mammalian cells^4^. Quiescence is characterized by a reversible arrest of cell proliferation, increased stress resistance, and reprogramming of gene expression and metabolism from a growth mode to a maintenance mode^9–12^. Quiescence is a highly prevalent yet under-studied state, with relevance for cellular and organismal aging. Quiescent states like dauer worms and non-dividing yeast cells have revealed universal aging-related processes, *e.g.* the conserved TORC1 nutrient-signalling network that controls quiescence entry and longevity from yeast to mammals by regulating growth, metabolism, and protein translation^4,12^. Human cells alternating between cellular quiescence and proliferation are critical for aging- and disease-associated processes, including stem-cell function, tissue homeostasis/renewal, immune responses, and drug resistance of tumors^11,13,14^. The aging and depletion of quiescent stem cells may be an important driver of organismal aging^13,15^.

Genomes are pervasively expressed, *e.g.* about 75% of the human genome is transcribed yet less than 2% codes for proteins. A substantial portion of transcriptomes consists of long non-coding RNAs (lncRNAs), which are longer than 200 nucleotides, do not overlap any coding RNAs, and can play varied roles in gene regulation at multiple levels^16–19^. Functional analyses of lncRNAs are challenging owing to poor annotation, low expression, and limited methodology^20–23^. Knowledge of lncRNAs is therefore far from complete even in well-studied organisms. Although lncRNAs show little sequence conservation, functional mechanisms and interacting proteins may be conserved^16,17^.

Mounting evidence implicates certain lncRNAs functioning in aging and associated diseases^24–31^. For example, lncRNAs play vital roles in aging-associated NFκB signalling^32^ and many lncRNAs are differentially expressed in aging human fibroblasts, *e.g.* lncRNA1 which delays senescence^33^. Certain lncRNAs are biomarkers for age-associated diseases and could provide more readily accessible drug targets than proteins^19,34,35^. Hence, lncRNAs are emerging as important, yet poorly characterized aging factors, presenting a promising research frontier. Fission yeast, featuring an RNA metabolism similar to metazoa, provides a powerful model system and living test tube to study lncRNA function^36–39^.

Recently, we have reported cellular phenotypes for 150 *S. pombe* lncRNA mutants in over 150 different nutrient, drug, and stress conditions^38^. Phenotype correlations revealed a cluster of four lncRNAs with roles in meiotic differentiation and the survival of quiescent spores. Here, we characterize one of these lncRNAs, *SPNCRNA.1530,* and show that it extends the CLS, interacts with ribosomal proteins, and reduces the ribosome content and cell growth. We name this lncRNA *aal1* for *aging-associated lncRNA1*. Remarkably, *aal1* expression in *Drosophila* is sufficient to extend the lifespan of flies, suggesting that the functional principle through which *aal1* acts is conserved in animals.

## Results

### *aal1* prolongs the chronological lifespan of non-dividing cells and reduces the growth of proliferating cells

The *aal1* gene (*SPNCRNA.1530*) encodes a 783 nucleotide lncRNA on Chromosome II. The *aal1* transcript levels are induced ∼6-24-fold in RNAi, nuclear-exosome, and other RNA-processing mutants (Figure 1A)^39^, suggesting that these RNA-processing pathways actively degrade *aal1* in proliferating cells. As is typical for lncRNAs, *aal1* is expressed below 1 RNA copy/cell on average in proliferating cells^9^, but its transcript levels are induced ∼6-30-fold in non-dividing cells, including stationary-phase cells, which are limited for glucose, as well as quiescent and meiotically differentiating cells, which are limited for nitrogen (Figure 1A). To validate these findings, we applied strand-specific RT-qPCR which showed that the *aal1* transcript levels are induced >15-fold in stationary-phase cells relative to proliferating cells (Figure 1B). The cycle threshold (Ct) after induction was 26.9±0.8, reflecting substantial *aal1* transcript levels in non-dividing cells.

**Figure 1:**
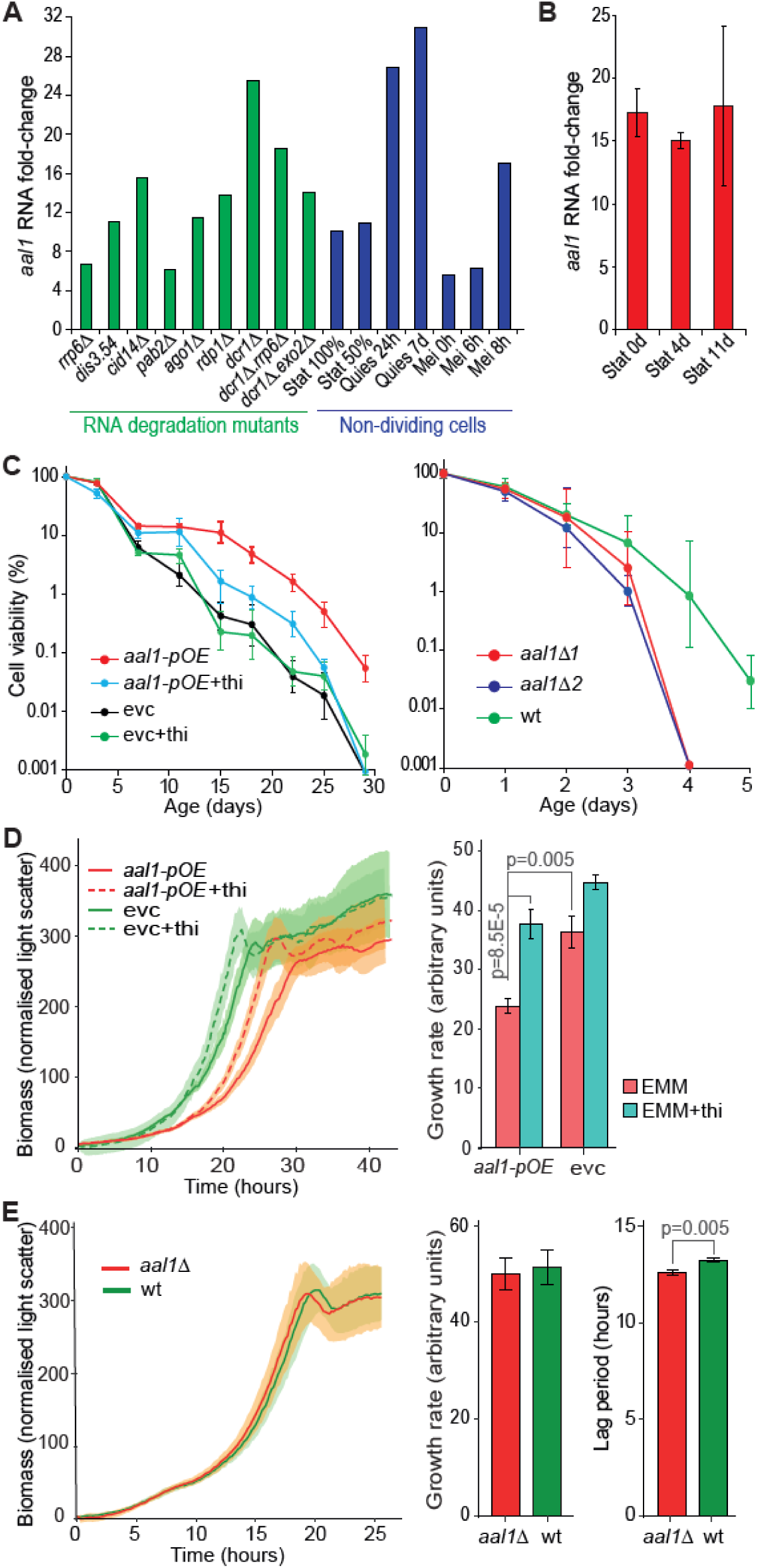
*aal1* prolongs the chronological lifespan of non-dividing cells and reduces growth of proliferating cells. (A) RNA fold-changes for *aal1* in different genetic and physiological conditions relative to wild-type proliferating cells, based on RNA-seq data^39^. Green: various RNA degradation mutants as indicated; blue: various non-dividing cells, including stationary-phase cells (Stat) at 100% and 50% viabilities, quiescent cells (Quies) after 24 hours and 7 days, and meiotically differentiating cells (Mei) at 0, 6, and 8 hours. (B) RNA fold-changes (log_2_) for *aal1* at the onset of stationary phase (Stat 0d) and after 4 and 11 days in stationary phase (Stat 4d and 11d) relative to proliferating cells, based on strand-specific RT-qPCR with gene-specific primers (ΔΔCt method). The *aal1* RNA levels are normalized to the lowly expressed coding gene *ppb1.* Bars indicate the standard errors (SE) of three independent repeats. (C) *Left graph*: Chronological lifespan assays for cells ectopically overexpressing *aal1* from a plasmid under the *P41nmt1* promoter (*aal1-pOE*) compared to empty-vector control cells (evc). Cells were cultured in the absence or presence of 15 µM thiamine (thi), where the *P41nmt1* promoter is active or repressed, respectively. *Right graph*: Chronological lifespan assays for *aal1* deletion cells (*aal1Δ*) compared to wild-type cells (wt). Two independent deletion strains (*aal1Δ1* and *aal1Δ2*) were generated by CRISPR-Cas9 using distinct sgRNAs. The percentages of viable cells were measured using a robotics-based colony-forming units (CFUs) assay^5^. A Poisson distribution-based model was used for maximum likelihood estimates of the number of CFUs and shown in the y-axes as percentage relative to the CFUs at Day 0. Data points reflect the mean ± SE of three biological repeats. Overexpression experiments which extend lifespan were performed in minimal medium, while deletion experiments which shorten lifespan were performed in rich medium. (D) *Left graph*: Ectopic *aal1* overexpression under the thiamine-repressible *P41nmt1* promoter (*aal1-pOE*) prolongs the lag period and reduces the growth rate compared to empty-vector control (evc) and/or in the absence of 15 µM thiamine (thi). Cells were grown in a microbioreactor and mean growth curves were fitted with *grofit* ^117^, with SD from six independent repeats shown as shades. *Right graph*: Quantitation of growth rate for experiments shown in the left graph. Growth rate calculations were done using *grofit*^117^ and statistical significance was determined with a one-way ANOVA followed by Tukey’s Honest Significant Difference test for pairwise comparisons in R^118^. Thiamine generally promotes growth in yeast cells^138^, as also observed here. (E) *Left graph*: Cells deleted for *aal1* (*aal1Δ*) show a slightly decreased lag period but similar growth rate to wild-type cells (wt). Same experimental setup and analysis as in (D). *Right graphs*: Quantitation of growth rates and lag periods for experiment in the left graph. Statistical significance was determined with one-way ANOVA followed by Dunnett’s test (*multcomp*)^119^ to correct for multiple testing of the comparisons of the lag periods and growth rates of *aal1Δ* against wt.

We tested the effects of *aal1* on the CLS of non-dividing cells. To this end, we constructed two strains, one for strong ectopic overexpression from a plasmid under the thiamine-repressible *P41nmt1* promoter (*aal1-pOE*) and one with a full deletion of the *aal1* gene (*aal1*Δ). Notably, *aal1-pOE* cells showed a prolonged lifespan during stationary phase, whereas *aal1*Δ cells showed a shortened lifespan (Figure 1C). Thus, *aal1* exerts an anti-aging, longevity effect in stationary phase cells.

The CLS is often inversely correlated with cell growth^6^. Therefore, we tested whether *aal1* affects the growth of proliferating cells. Indeed, *aal1-pOE* cells featured longer lag periods and slower growth rates compared to control cells (Figure 1D). Conversely, *aal1Δ* cells showed only a subtly reduced lag period (Figure 1E). This weak positive growth effect in proliferating *aal1Δ* cells is consistent with *aal1* being hardly expressed in these cells anyway^9^. We conclude that *aal1* has an anti-growth effect in proliferating cells.

While the *aal1* gene locus does not overlap any protein-coding genes, another lncRNA (*SPNCRNA.401*) is located antisense to *aal1* with over 90% sequence overlap (Supplemental Figure 1A). This setup renders it impossible to distinguish between deletions of *aal1* and *SPNCRNA.401,* raising the possibility that phenotypes observed in *aal1*Δ cells could be caused by the absence of *SPNCRNA.401.* Like *aal1, SPNCRNA.401* is induced in non-dividing cells^39^. We, therefore, checked whether overexpression of *SPNCRNA.401* might show effects on lifespan and growth similar to overexpression of *aal1*. As for *aal1,* we constructed a strain for ectopic overexpression (*SPNCRNA.401-pOE*). In contrast to *aal1-pOE* cells, the *SPNCRNA.401-pOE* cells did not show any significant effects on lifespan or cell growth (Supplemental Figure 1B). If anything, *SPNCRNA.401-pOE* cells showed inverse, but much weaker phenotypes than *aal1-pOE* cells, by marginally decreasing lifespan but increasing cell growth. Moreover, ectopic overexpression of *aal1* was sufficient to rescue the short lifespan of *aal1Δ* cells, further supporting that the CLS phenotype reflects the absence of *aal1* (Supplemental Figure 1C). Given that *aal1* is close to the promoters of the flanking coding genes *ckb2* and *ctu1* (Supplemental Figure 1A) and some lncRNAs regulate the expression of neighboring genes *in cis*^40–42^, we also quantified the changes in transcript levels of these neighboring genes by qRT-PCR. These experiments showed that deletion of *aal1* did at most marginally affect the expression of the neighboring genes (Supplemental Figure 1D). Taken together, we conclude that the phenotypes observed in *aal1Δ* and *aal1-pOE* cells reflect the absence and overexpression, respectively, of the *aal1* RNA.

Inhibition of the conserved Target of Rapamycin Complex 1 (TORC1) signaling network leads to decreased cell growth and prolonged lifespan from yeast to mammals^43,44^. The *aal1* overexpression phenotypes resemble those of TORC1 inhibition, raising the possibility that *aal1* functions within the TORC1 network. Therefore, we tested whether the *aal1* growth phenotype depends on TORC1 function. TORC1 inhibition by rapamycin and caffeine led to similar relative growth reduction in *aal1-pOE* and *aal1Δ* cells as in the respective controls (Supplemental Figure 2). These results indicate that overexpression of *aal1* and inhibition of TORC1 exert additive effects on cell growth, suggesting that *aal1* acts independently of the TORC1 network.

We conclude that the expression of the *aal1* lncRNA strongly increases in non-dividing cells, and the absence or excess of the *aal1* RNA is sufficient to shorten or extend the lifespan of these cells, respectively. Moreover, expression of *aal1* in proliferating cells leads to reduced growth. The lifespan and growth phenotypes are mediated by *aal1* independently of the TORC1 network. Notably, *aal1* can promote longevity and repress growth when expressed from a plasmid, indicating that it acts *in trans* as a lncRNA.

### *aal1* genetically interacts with coding genes functioning in protein translation and localises to the cytoplasm

To obtain clues on the molecular function of *aal1*, we systematically assayed genetic interactions between the non-coding *aal1* gene and protein-coding genes. Combining two mutations in the same cell can cause phenotypes that are more or less severe than expected from the phenotypes of the single mutations, defining negative and positive genetic interactions, respectively. Such genetic (or epistatic) interactions reveal broad relationships between functional modules or biological processes^45^. To systematically uncover functional relationships between the *aal1* RNA and proteins, we used an *aal1Δ* strain (*aal1::natMX6*) as the query mutant to screen for interactions with all 3420 non-essential coding-gene deletions with the synthetic genetic array (SGA) method^46,47^. We measured all pairwise genetic interactions using colony size as a proxy for double-mutant fitness relative to *ade6Δ* control double mutants. As for screens with coding-gene query mutants^6,48^, we observed moderate reproducibility between the three repeats and more negative than positive interactions, revealing 140 negative and 28 positive genetic interactions, respectively, for at least two out of the three repeats (Supplemental Dataset 1). Analysis using AnGeLi^49^ showed that these interacting genes were enriched in those showing lifespan and growth phenotypes, as *aal1* itself, including the fission yeast phenotype ontology (FYPO) terms^5,8,50^ ‘loss of viability in stationary phase’ (*p_adj_* = 5.3E-17) and ‘abnormal vegetative cell population growth’ (*p_adj_* = 5.0E-14). Moreover, the interacting genes were strongly enriched for several Gene Ontology (GO) terms^51^ related to ribosome biogenesis/function and cytoplasmic translation (Figure 2A). We conclude that genetic interactions for *aal1* point to functions associated with protein translation.

**Figure 2:**
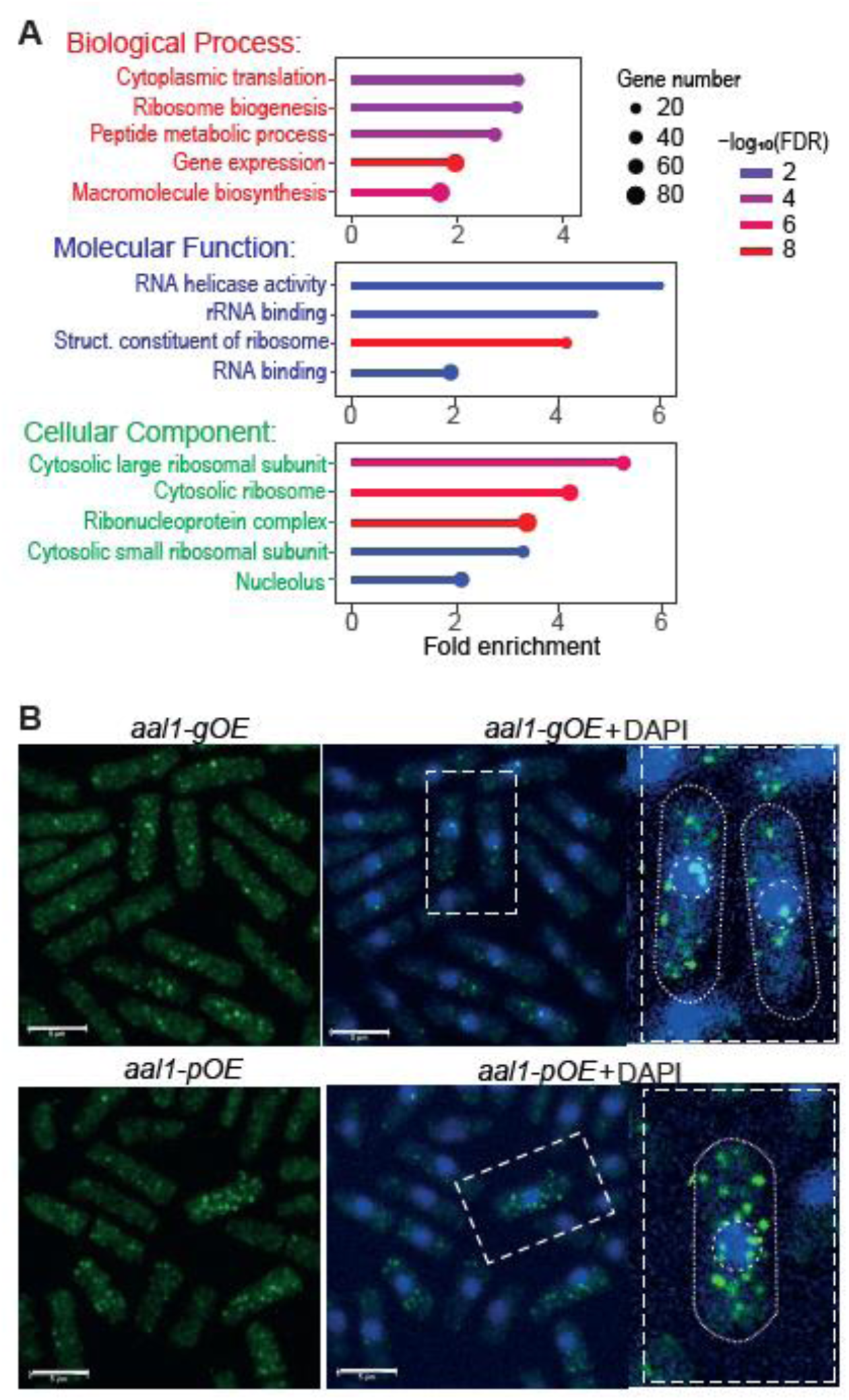
*aal1* shows functional relationships with translation-related processes and localizes to the cytoplasm. **(A)** GO-term enriched among the genes that showed positive or negative genetic interactions (FDR ≤0.05) in at least 2 of the 3 repeats in the SGA screens using *aal1Δ* as query mutant. Representative GO terms for Biological Process (red), Molecular Function (blue), and Cellular Component (green) are shown, selected for non-redundancy, specificity, and significance. The graphs show the fold enrichments as well as the gene numbers and −log_10_ of false-discovery rate as indicated in the legends at right. Visualisation with ShinyGO (ver 0.77)^139^. The genetic-interaction and background gene lists are provided in Supplemental Dataset 1. **(B)** Fluorescence micrographs of single-molecule FISH experiments of *aal1-gOE* cells (*top*) and *aal1-pOE* cells (*bottom*). The *aal1* RNAs are labelled in green and DAPI-stained DNA is shown in blue. Enlarged cells with nuclei are outlined at right. Scale bars: 5 μm.

To further test the possibility that *aal1* functions in translation, we analyzed the subcellular localization of the *aal1* RNA using antisense probes for single-molecule RNA fluorescence *in situ* hybridization (smRNA-FISH)^52^. Given that the *aal1* RNA expression is very low in proliferating cells (Figure 1A,B) and smRNA-FISH is challenging in stationary-phase cells^53^, we generated a strain expressing *aal1* from the *P41nmt1* promoter at its native genomic locus (*aal1-gOE*). This analysis revealed that *aal1* RNAs predominantly localize in the cytoplasm in multiple foci (Figure 2B). Similar results were obtained with the *aal1-pOE* strain that ectopically overexpresses *aal1* (Figure 2B). This finding corroborates that *aal1* functions *in trans* as a lncRNA and is consistent with a role of *aal1* in cytoplasmic translation.

### *aal1* associates with ribosomes and reduces the cellular ribosome content

Many lncRNAs function with RNA-binding proteins (RBPs), and identifying such RBPs can shed light on the molecular roles of the target lncRNAs^54^. To uncover RBPs that interact with *aal1*, we applied an RNA-centric approach, ‘comprehensive identification of RBPs by mass spectrometry’ (ChIRP-MS)^55^. We designed biotinylated antisense oligos along the length of *aal1* to pull down RBPs associated with *aal1*, which does not require genetic manipulations that could interfere with *aal1* function (Methods). We performed ChIRP-MS with wild-type and *aal1-pOE* cells after 6 days in stationary-phase, when the *aal1* transcript levels are induced, with *aal1Δ* cells serving as control. In total, we identified 218 proteins across all three independent repeats in all strains, 68 of which were significantly more abundant in the pull-downs from wild-type and/or *aal1-pOE* cells than from *aal1Δ* cells (FDR ≤0.005 and log_2_FC ≥6.5; Supplemental Dataset 2). These 68 *aal1*-bound proteins were enriched for biological processes associated with cytoplasmic translation and energy metabolism (Figure 3A). The 68 proteins were more highly connected with each other than expected by chance for known protein-protein interactions, reflecting a large network of ribosomal and ribosome-associated proteins (Supplemental Figure 3). While 27 of the 68 proteins have known functions in translation-related processes, most of the remaining proteins are enzymes functioning in energy metabolism. Recent research indicates that many metabolic enzymes exert moonlighting functions as RBPs that can act as ribosome-associated proteins^56,57^. Notably, 44 of the 68 *aal1*-bound proteins have been independently shown to be RBPs in *S. pombe* (Supplemental Dataset 2)^57^. Taken together, these results indicate that *aal1* directly or indirectly binds to multiple proteins associated with ribosomes and/or ribosomal subunits.

**Figure 3:**
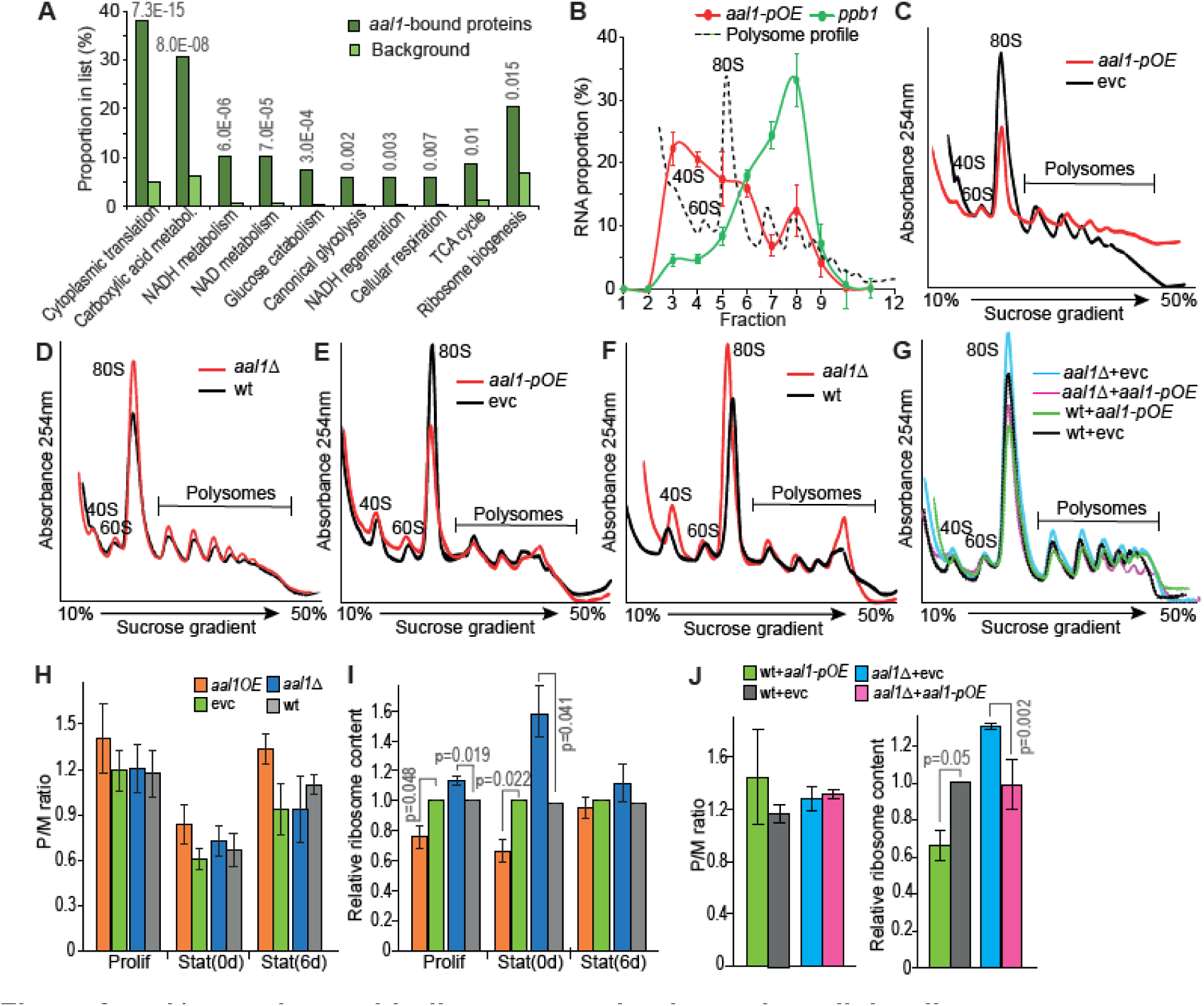
*aal1* associates with ribosomes and reduces the cellular ribosome content. **(A)** Graph showing the proportions of genes with representative GO Biological Process terms among the 68 *aal1*-bound proteins compared to all *S. pombe* proteins (background). GO terms were selected for non-redundancy, specificity and significance, with the respective enrichment FDRs shown on top. The gene list used is provided in Supplemental Dataset 2. **(B)** Polysome fractionation followed by RT-qPCR for *aal1-pOE* cells shows that *aal1* (red curve) mainly occurs with free ribosomal subunits (40S/60S) and monosomes (80S). The *ppb1* control mRNA (green curve) mainly occurs in polysomes. The corresponding polysome profile is shown as a black, dashed line. Enrichment was calculated relative to the free RNA Fractions 1 and 2^59^. The plot shows the mean ± SE of three independent repeats of *aal1* overexpressing cells during exponential growth in minimal medium. **(C)** Polysome profiling for proliferating *aal1-pOE* and empty-vector control (evc) cells in minimal medium. The two profiles are aligned at the lowest points of the monosome peaks, corresponding to the baseline. The polysome profiles from two independent biological repeats are shown in Supplemental Figure 4B. **(D)** Polysome profiling for proliferating *aal1Δ* and wild-type cells as in (C). The polysome profiles from two independent biological repeats are shown in Supplemental Figure 4B. **(E)** Polysome profiling for late stationary-phase (Day 6) *aal1-pOE* and evc cells as in (C). The polysome profiles from two independent biological repeats are shown in Supplemental Figure 4B. **(F)** Polysome profiling for late stationary-phase (Day 6) *aal1Δ* and wild-type cells as in (C). The polysome profiles from two independent biological repeats are shown in Supplemental Figure 4B. **(G)** Polysome profiling for proliferating cells in minimal medium. Ectopic expression of *aal1* in *aal1Δ* background (*aal1Δ* + *aal1-pOE*) reduces the increased ribosome content observed in *aal1Δ* cells to a content similar to wild-type cells expressing an empty vector (wt+evc). The polysome profiles from two independent biological repeats are shown in Supplemental Figure 4C. **(H)** Polysome/monosome (P/M) ratio in proliferating cells (Prolif), at the onset of stationary phase, Stat(0d), and after 6 days in stationary phase, Stat(6d), in *aal1-pOE*, empty-vector control (evc), *aal1Δ* and wild type (wt) cells as indicated. Bars show mean ± SE of three independent repeats, with statistical significance determined using two-sample t-test. **(I)** Relative ribosome content (Polysome+Monosome) in proliferating cells (Prolif), at the onset of stationary phase, Stat(0d), and after 6 days in stationary phase, Stat(6d), in *aal1-pOE*, relative to empty-vector control (evc) and *aal1Δ* relative to wild type (wt) cells as indicated. Bars show mean ± SE of three independent repeats, with statistical significance determined using two-sample t-test against the respective controls. The differences at Stat(6d) are not significant, likely reflecting that the ribosome content is generally low at this stage which makes it harder to obtain reproducible measurements. **(J)** Polysome/monosome (P/M) ratios and ribosome content as in (H) and (I) for ectopic expression experiments shown in (G). Relative ribosome content determined relative to wt+evc.

To corroborate and elucidate this ribosomal association, we applied polysome fractionation^58^ followed by RT-qPCR analysis^59^. Most *aal1* RNA was detected together with free ribosomal subunits (40S/60S) and monosomes (80S) in proliferating *aal1-pOE* cells (Figure 3B). As a control mRNA, we used *ppb1,* which is the least variable gene under many perturbations, including stationary phase, and is comparatively lowly expressed^60^. As expected, *ppb1* was predominantly found in polysomes (Figure 3B). Similar *aal1* localization patterns were apparent for both wild-type and *aal1-pOE* cells in young and old stationary-phase cells, although in the older cells, a substantial proportion of *aal1* was also detected in polysomes (Supplemental Figure 4A).

We then checked for any effects of *aal1* on the polysome profiles. Remarkably, proliferating *aal1-pOE* cells showed a decreased polysome profile compared to empty-vector control cells (Figure 3C), while proliferating *aal1Δ* cells showed an increased profile compared to wild-type control cells (Figure 3D). These differences were most pronounced for the monosome fractions. Similar patterns were evident in the polysome profiles from *aal1-pOE* and *aal1Δ* cells during early and late stationary phase (Figure 3E,F; Supplemental Figure 4B). These results suggest that *aal1* has an inhibitory effect on the cellular ribosome content. To establish that the increased ribosome content in *aal1Δ* cells was caused by the absence of *aal1*, we ectopically expressed *aal1* (*aal1-pOE*) or an empty-vector control in *aal1Δ* cells. Indeed, ectopic expression of *aal1* was sufficient to decrease the ribosome content in *aal1Δ* cells to levels similar to or below those in wild-type cells (Figure 3G; Supplemental Figure 4C). Together, these results indicate that *aal1* can act *in trans* to reduce the cellular ribosome content, while the absence of *aal1* increases the ribosome content.

To quantify and validate the observed differences in the polysome profiles, we measured the polysome/monosome ratios (P/M; estimate of translational efficiency) and ribosome content (P+M; estimate of translational capacity) for all polysome profiles of *aal1-pOE, aal1Δ* and control cells (Figure 3C-F; Supplemental Figure 4B). The P/M ratios did not show significant changes in *aal1Δ* or *aal1-pOE* cells (Figure 3H). However, *aal1* overexpression and deletion led to decreased and increased cellular ribosome contents, respectively, relative to the respective controls, although these changes were only significant in proliferating and early stationary phase cells (Figure 3I). We also quantified these parameters in the ectopic expression experiments in Figure 3G and Supplemental Figure 4C. Again, the differences were larger for ribosome content than for P/M ratios, with ectopic overexpression of *aal1* significantly reducing the ribosome content of *aal1Δ* cells (Figure 3J). Thus, this analysis confirms the *aal1*-mediated differences in ribosome content. Taken together, these findings indicate that *aal1* associates with ribosomes and that it can reduce the cellular content of ribosomes.

### *aal1* binds to the *rpl1901* mRNA encoding a ribosomal protein whose repression prolongs lifespan

To further dissect how *aal1* might affect the cellular ribosome content, we used RNA-seq to examine its effects on the transcriptome. We analysed the differential gene expression between *aal1-pOE* and empty-vector control cells after 6 days in stationary phase, revealing 79 induced and 248 repressed transcripts. The overexpression of *aal1* led to the repression of ribosome- and translation-related transcripts, including 127 of 135 mRNAs encoding *S. pombe* ribosomal proteins (Figure 4A; Supplemental Dataset 3). The induced genes, on the other hand, were not enriched in any biological processes. Thus, *aal1* has the capacity to directly or indirectly down-regulate the mRNAs for key proteins involved in translation.

**Figure 4:**
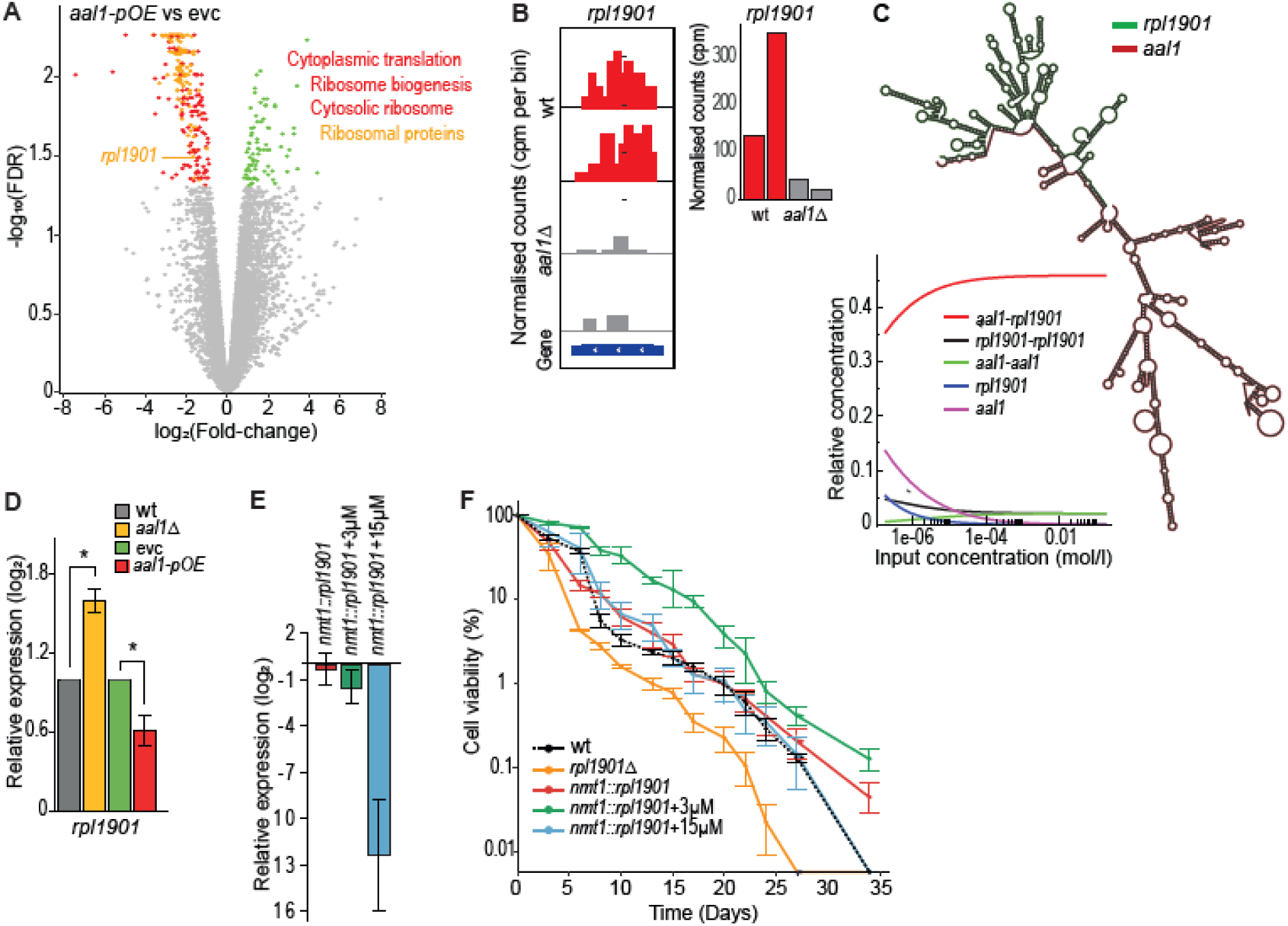
*aal1* binds to the *rpl1901* mRNA whose repression prolongs lifespan. **(A)** Volcano plot of RNA-seq data comparing transcript levels in *aal1-pOE* relative to empty-vector control cells during stationary phase (Day 6) in minimal medium. The 248 repressed genes (red) are enriched in the GO terms ‘Cytoplasmic translation’ (139 genes; FDR 5.7E-135), ‘Ribosome biogenesis’ (74 genes; 8.6E-32), and ‘Cytosolic ribosome’ (134 genes; FDR 9.2E-178). The 127 repressed ribosomal protein genes are highlighted in orange with *rpl1901* indicated. The 79 induced genes (green) are not enriched in any GO terms. **(B)** The ribosomal-protein mRNA *rpl1901* is enriched in two independent repeats of ChIRP pull-downs from wild-type (wt, red) compared to *aal1Δ* (grey) cells. *Left panels*: strand-specific ChIRP-seq reads in counts per million (cpm) for 50 bp bins across *rpl1901,* visualized using IGV tracks^139^ and deepTools^134^. *Right panel*: normalised read counts (cpm) for *rpl1901*. **(C)** *In silico* predictions of *aal1-rpl1901* interaction using the ViennaRNA package^135^ with RNAcofold^136^. *Top*: Predicted *aal1*-*rpl1901* heterodimer structure showing the interaction sites along with potential RNA secondary structures. *Bottom*: Concentration dependency plot of dimerization showing the computed homo- and hetero-dimerizations of RNAs for concentration relative to each other (y-axis) and different input concentrations (x-axis), with predicted equilibrium concentrations for the two monomers, *aal1* and *rpl1901,* the two homodimers, *aal1-aal1* and *rpl1901-rpl1901,* and the *aal1-rpl1901* heterodimer. **(D)** RT-qPCR experiment to quantify RNA levels of *rpl1901* in proliferating wild-type (wt), *aal1Δ*, empty-vector control (evc), and *aal1-pOE* cells. Expression was normalised to *act1* and shown relative to the expression in the respective controls. Bars show the mean ± SE of three replicates; asterisks indicate p-values ≤0.05, determined by t-test. **(E)** RT-qPCR experiment to quantify expression of *rpl1901* under the thiamine-repressible *P41nmt1* promoter at its native locus (*nmt1::rpl1901*), in the absence or presence of 3 or 15 µM thiamine as indicated. Expression was normalised to *act1* and shown relative to wild-type expression. Bars show the mean ± SE of three replicates. **(F)** Chronological lifespan assays for cells expressing *rpl1901* at different levels. Only the moderate repression of *rpl1901* with 3 µM thiamine promotes lifespan extension. Experiments were performed in minimal medium. Assays and analyses as in Figure 1C.

Some lncRNAs can regulate genes post-transcriptionally, *e.g.* through partial base-pairing between a lncRNA and target mRNAs to promote mRNA degradation^61^ or inhibit translation^62^. To test whether *aal1* binds to other RNAs, we used the pull-downs from the ChIRP-MS experiment to sequence any RNAs associated with *aal1* (ChIRP-seq)^63^. As expected, *aal1* itself was the top-enriched RNA in these pull-downs compared to *aal1Δ* control cells, but other RNAs, both coding and non-coding, also appeared to be enriched (Supplemental Figure 5A; Supplemental Dataset 4). Among these RNAs were four encoding ribosomal proteins, of which *rpl1901* was most consistently enriched (Figure 4B). To test the plausibility of an RNA-RNA interaction between *aal1* and *rpl1901*, we performed an *in silico* analysis using the ViennaRNA Package. This analysis predicted that *aal1* has a high potential to dimerize with *rpl1901* at very low concentrations (Figure 4C), thus supporting the interaction between these two RNAs.

As with most other ribosomal-protein genes, *rpl1901* was repressed upon *aal1* overexpression in chronologically aging cells (Figure 4A). To quantitatively analyze how *aal1* affects *rpl1901* mRNA levels, we performed RT-qPCR assays in both *aal1Δ* and *aal1-pOE* cells during proliferation. RNA levels of *rpl1901* were modestly (∼2-fold) but significantly repressed in *aal1-pOE* cells and induced in *aal1Δ* cells (Figure 4D). This result is consistent with the possibility that *aal1* directly represses the *rpl1901* mRNA levels, which could contribute to the *aal1*-mediated reduction in ribosome content and lifespan extension.

We wondered whether repression of *rpl1901* might be involved in the *aal1-*mediated lifespan extension. We first generated a deletion mutant of *rpl1901 (rpl1901Δ*) and showed that *rpl1901Δ* cells featured substantially shorter CLS and slower growth than wild-type cells (Supplemental Figure 5B). However, *rpl1901* was completely absent in this deletion mutant, whereas *rpl1901* expression was only moderately repressed in response to *aal1* (Figure 4D). Therefore, we tested whether a moderate repression of *rpl1901* might extend the lifespan. To this end, we expressed *rpl1901* under the thiamine-repressible *P41nmt1* promoter in its native locus (*nmt1::rpl1901*) to modulate *aal1* repression by supplementing various doses of thiamine. The expression levels of *nmt1::rpl1901* in the absence of thiamine were comparable to the *rpl1901* levels under its native promoter (Figure 4E). Addition of 3 µM thiamine led to a ∼2-fold reduction in *nmt1::rpl1901* levels similar to the repression of *rpl1901* observed in *aal1-pOE* cells, while 15 µM thiamine led to a much stronger repression (Figure 4D,E). Expression of *nmt1::rpl1901* prolonged the lag period but not the growth rate, while in the presence of 3 or 15 µM thiamine, the growth rates were modestly but significantly reduced (Supplemental Figure 5C). Notably, only the ∼2-fold repression of *rpl1901* with 3 µM thiamine led to a substantial extension of the CLS, while the other conditions, leading to no or much stronger repression of *rpl1901*, showed similar or shorter CLS compared to wild-type cells (Figure 4F). We conclude that only a moderate (∼2-fold) repression of *rpl1901* but not a stronger repression is beneficial for cellular longevity, mirroring the *rpl1901* repression and lifespan extension seen in *aal1-pOE* cells.

### *aal1* can extend the lifespan of flies

Ribosomal proteins and most of the translational machinery are highly conserved across eukaryotes, and reducing protein translation to non-pathological levels prolongs lifespan in different model organisms^64–67^. Although lncRNAs show little or no sequence conservation between species, their functional principles can be conserved^16,17^. We wondered whether *aal1* might have the potential to extend lifespan in a multicellular eukaryote, as it does in *S. pombe* cells. We tested this possibility in the fruit fly, *Drosophila melanogaster*. We first ensured the expression of *aal1* in *Drosophila* by confirming the presence of the RNA of the expected size in female flies where *UAS-aal1* was driven by the ubiquitous, constitutive *GAL4* driver *daughterlessGAL4* (Supplemental Figure 6A). We further observed that such ubiquitous expression of *aal1* in *Drosophila* throughout development resulted in a significant reduction in the number of flies that reached adulthood compared to control flies, indicating that *aal1* expression negatively affects development (Supplemental Figure 6B). To assess the impact of *aal1* on *Drosophila* aging, we focused on the gut because *aal1* expression in yeast represses protein translation, and interventions that repress translation in the fly gut have pro-longevity effects^68,69^. To avoid any deleterious effects on development, we used the mid-gut specific RU486 inducible driver (*TIGS*) in adult females which can drive *aal1* expression upon feeding the inducer, RU486. Remarkably, this induction of *aal1* significantly extended the medium lifespan of female flies (Figure 5A). The feeding of the inducer itself did not affect the lifespan in the *TIGS* driver-alone control (Figure 5B), indicating that the extension is not an artifact of RU468 feeding. Although *aal1* is not conserved in *Drosophila,* it has a high predicted potential to dimerize with the *RpL19* mRNA, encoding the highly conserved *Drosophila* ortholog of Rpl1901 (Supplemental Figure 6C). These findings show that *aal1* can promote longevity in *Drosophila* and raise the possibility that the functional principle of *aal1* is conserved and that *aal1* lncRNA counterparts might exist in multicellular eukaryotes.

**Figure 5:**
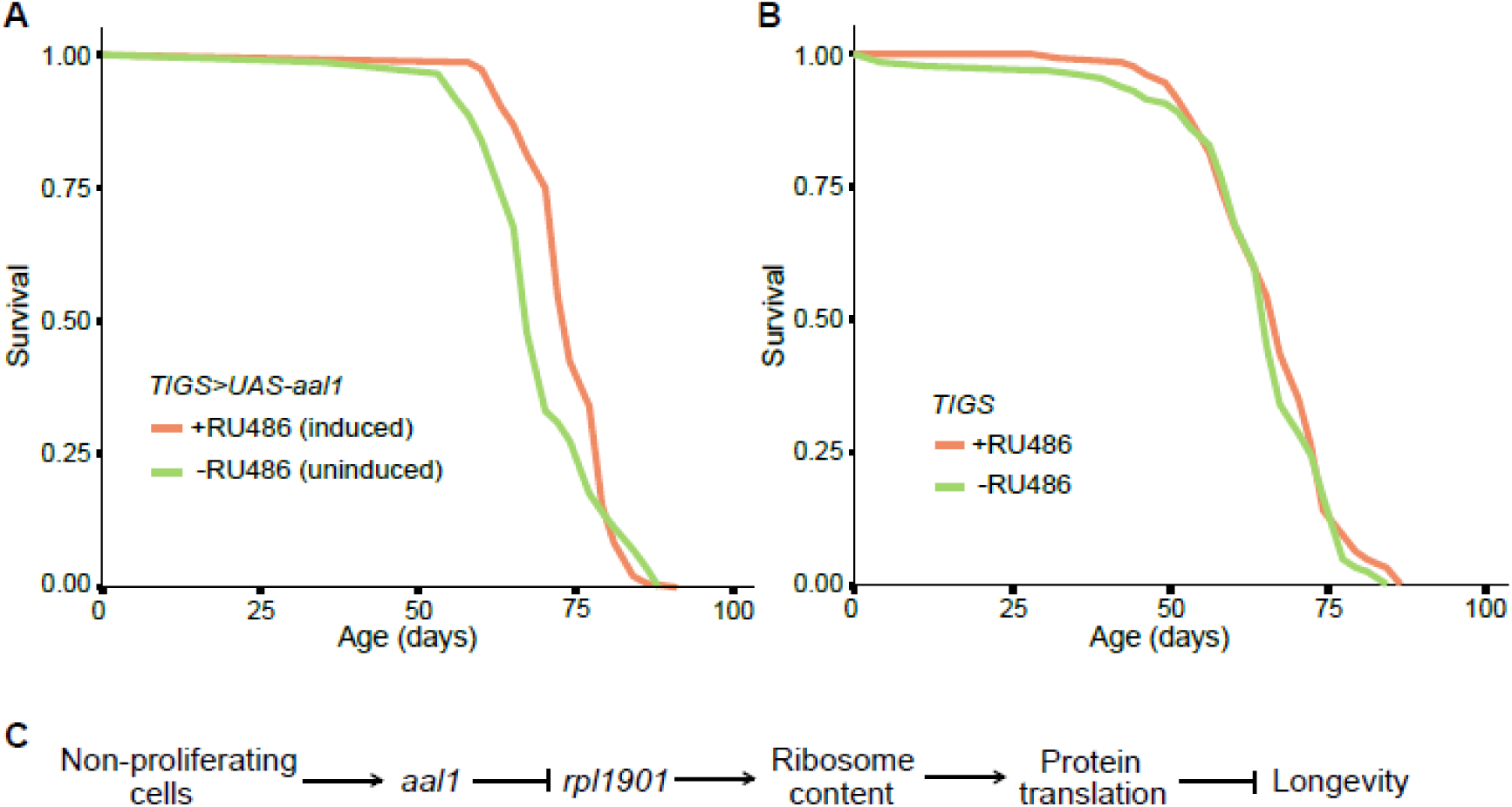
*aal1* expression prolongs the lifespan in flies. (A) Lifespans of female *Drosophila* with *UAS-aal1* induced in adult midguts with the gut-specific *TIGS* driver by RU486 feeding to activate expression (−RU486: n = 142 dead/8 censored flies, +RU486: n = 144 dead/6 censored flies, p = 0.003, log-rank test). (B) Lifespans of control females carrying the *TIGS* driver alone with or without RU486 feeding (−RU486: n = 131 dead/19 censored flies, +RU486: n = 144 dead/6 censored flies, p = 0.3, log-rank test). (C) Simple model for *aal1* function in lifespan extension via inhibition of *rpl1901* expression and protein translation (details in Discussion).

## Discussion

This study presents the functional characterization of a lncRNA, *aal1*, which can lower the ribosomal content and extend the lifespan of non-dividing fission yeast cells. The genetic and protein interactions of *aal1*, along with its various phenotypes, point to an involvement in protein translation. The *aal1* RNA localizes to the cytoplasm and its ectopic expression from a plasmid can rescue *aal1* deletion phenotypes and trigger key phenotypes like decreased ribosome content, reduced growth, and increased lifespan, indicating that *aal1* functions *in trans*. While further work is required to dissect the specific mechanisms of *aal1* function, we propose a simple model for how *aal1* might exert its roles in protein translation and longevity (Figure 5C). Below, we discuss the key steps of this model in the context of our results and published findings.

The *aal1* RNA is induced in non-dividing cells, possibly by inhibiting its degradation via the RNAi and nuclear-exosome pathways that repress *aal1* in proliferating cells^39^ (Figure 1A,B). Then, *aal1* associates with other RNAs, including the *rpl1901* mRNA that encodes a ribosomal protein (Figure 4B,C). This interaction may lead to a ∼2-fold reduction of *rpl1901* mRNA levels (Figure 4D), and such a modest reduction is critical and sufficient to extend the CLS (Figure 4F). Overexpression of *aal1* leads to a similar *rpl1901* reduction (Figure 4D) and CLS extension (Figure 1C).

How might *aal1* repress the *rpl1901* mRNA levels? It is known that lncRNAs can regulate genes post-transcriptionally through base-pairing with target mRNAs to promote their degradation or inhibit their translation^61,62^. The *aal1* RNA associates with ribosomes (Figure 3A,B; Supplemental Figure 3), as also reflected in the genetic interactions (Figure 2A), pointing to its mode of action. Several RNAs interact with ribosomes in different organisms^70–73^, reflecting their translation into peptides^72,73^, degradation via nonsense-mediated decay (NMD)^74^, or involvement in translational control^62,71,75^. Available data indicate that *aal1* is not detectably translated in cells undergoing meiosis^72^, when *aal1* is induced (Figure 1A), and *aal1* is not induced in mutants of the NMD-pathway gene *upf1*^39^. It is possible, however, that the interaction of *aal1* with the *rpl1901* mRNA during its translation leads to NMD-mediated degradation of *rpl1901.* We find that *aal1* is more associated with monosomes than polysomes (Figure 3B), which might suggest a role before translational elongation, although older cells also showed a substantial proportion of *aal1* in polysomes (Supplemental Figure 4A). Notably, monosomes can also reflect the active translation of short open reading frames (<590 nt), where translational initiation takes relatively more time than translational elongation^76,77^. The *rpl1901* mRNA is 582 nt long, so our results are consistent with the possibility that *aal1* associates with *rpl1901* in the context of translating ribosomes. Other or additional pathways may contribute to an *aal1*-mediated *rpl1901* mRNA repression. However, RNAi does not seem to be directly involved in this repression, because the *rpl1901* mRNA, unlike *aal1* (Figure 1A), is reduced in RNAi mutants^39^.

Compared to *rpl1901, aal1* is more lowly expressed, raising a potential paradox of regulatory stoichiometry. But in non-dividing cells, *rpl1901* is expressed at only ∼6 molecules per cell^9^, which may be similar to the *aal1* which is induced in this condition (Figure 1A). Furthermore, several lncRNAs, including Xist and NORAD, form biomolecular condensates to facilitate their substoichiometric action, as these phase-separated condensates can concentrate factors^78,79^. Indeed, the low levels of the Xist RNA relative to its abundant targets is essential to maintain target specificity, suggesting that low levels are critical for lncRNA function^80,81^. Examples of ribonucleoprotein condensates are stress granules, processing bodies, and the nucleolus^82^. As an example in *S. pombe, meiRNA*, a lncRNA expressed at only 0.21 molecules per cell^9^, forms a nuclear body with the RNA-binding protein Mei2 to promote meiotic differentiation^83^. Recently discovered translation factories are cytoplasmic bodies accumulating many copies of certain RNAs^84^. Notably, we observed that *aal1* localizes in cytoplasmic foci (Figure 2B), which might reflect ribonucleoprotein bodies facilitating molecular interactions between *aal1, rpl1901,* and ribosomes to trigger NMD-mediated degradation of *rpl1901*.

We propose that the *aal1-*mediated repression of *rpl1901* then leads to the global repression of other ribosomal protein genes, which in turn reduces the cellular ribosome content and protein translation, thus promoting longevity (Figure 5C). Gene-specific feedback mechanisms can increase the expression of individual ribosomal proteins present at substoichiometric amounts, but these mainly function through the regulation of splicing and/or mRNA stability rather than transcription or translation^85^. The *rpl1901* gene does not have introns and *aal1-*mediated degradation might lead to an attenuation of protein translation. This could be achieved by global negative feedback regulation of other ribosomal protein genes in response to lowered *rpl1901* levels^86,87^. Consistent with this possibility, a ∼2-fold reduction in *rpl1901* levels is sufficient to reduce cell growth (Supplemental Figure 5C). Moreover, *aal1* overexpression leads to the repression of 127 of 135 mRNAs encoding the proteins of the *S. pombe* cytoplasmic ribosome (Figure 4A) and triggers a reduction in cellular ribosome content (Figure 3) and growth (Figure 1D). Ribosome content, translational capacity, and growth rate are linked in yeast and other cells, *e.g.* via TORC1 signaling^44,87–90^. As would be expected for the proposed mechanism, however, the *aal1-*mediated growth reduction is independent of TORC1 signaling (Supplemental Figure 2). Attenuation of protein translation to non-pathological levels is known to prolong lifespan and delay aging-associated diseases in model organisms, although the detailed nature of these relationships is still unclear^64–67,91–93^.

It is of course possible that *aal1* functions via additional or other processes than those proposed in Figure 5C. However, it is notable that modulation of *rpl1901* expression is sufficient to mirror the key phenotypes of *aal1.* Intriguingly, while mutants in most ribosomal proteins do not appear to affect aging^94^, orthologs of Rpl1901 (RPL19) are among only four ribosomal proteins shown to affect aging in budding yeast and worms where the loss of RPL19 function extends lifespan^95–97^. RPL19 is also among the few ribosomal protein genes that are up-regulated in embryonic stem cells of human blastocysts and hepatocellular carcinoma^98,99^, and post-transcriptional silencing of RPL19 alleviates human prostate cancer by controlling the translation of a subset of transcripts^100^. Ribosomal proteins may affect aging in different ways, *e.g.* by altering global translation rates, functioning in specialized ribosomes, or affecting the protein biogenesis machinery^101,102–104^.

Protein translation is highly energy-demanding and tightly regulated by conserved mechanisms to adjust supply with demand in varying environmental or physiological conditions^66^. Non-coding RNAs, including the well-known rRNAs, tRNAs, snoRNAs and miRNAs, function in protein translation and its regulation. More recently, lncRNAs have emerged as regulators of ribosome biogenesis and translation. For example, SLERT controls nucleolar phase separation and rRNA expression^105^, LoNA inhibits translation and rRNA expression at multiple levels^106^, while other lncRNAs promote rDNA chromatin compaction to repress rRNA transcription in quiescent cells^107^. Several ribosome-associated non-coding RNAs are emerging as translational regulators in response to stress^108,109^. The nematode *daf-2* mutants in an insulin receptor-like gene display similar phenotypes to *aal1,* including slow growth and extended lifespan^110^, along with low ribosomal-protein levels and reduced protein synthesis^111,112^. Interestingly, the *tts-1* RNA is required for the lifespan extension in *daf-2* mutants and is mostly associated with the monosomal fraction of ribosomes^75^, akin to *aal1*, raising the possibility that these lncRNAs play similar mechanistic roles.

To our surprise, the expression of *S. pombe aal1* in *Drosophila* leads to a significant lifespan extension in *Drosophila* (Figure 5A). Interestingly, both the organ whence the longevity effects arise, namely the gut, and the magnitude of the effects are comparable between *aal1* expression and the partial inhibition of RNA polymerase I, which transcribes the pre-rRNA^69^. Thus, although *aal1* is not conserved in *Drosophila,* this RNA can substantially promote the longevity of both single yeast cells and multicellular flies. Furthermore, given that ribosomal proteins and translational processes are highly conserved from yeast to humans, it is plausible that *aal1* functions through related mechanisms in both yeast and flies, and perhaps can take on the role of a native fly lncRNA which functions within a conserved regulatory framework. Indeed, *in silico* analysis predicts that *aal1* has a high potential to dimerize with the *Drosophila RpL19* mRNA, encoding the highly conserved ortholog of *S. pombe* Rpl1901 (Supplemental Figure 6C). Structure-function constraints for lncRNAs are more relaxed than for proteins^74^, and RNAs with different sequences may interact with similar proteins. It would be interesting to test whether *aal1* binds the *RpL19* mRNA in *Drosophila* to inhibit the cellular ribosome content. The finding in *Drosophila* raises the intriguing possibility that other organisms use analogous lncRNAs and related mechanisms to control the translational capacity of certain cells, and such lncRNAs would potentially be effective targets for future RNA-based therapies to delay aging and associated diseases.

## Methods

### Fission yeast strains and culture conditions

The standard *S. pombe* lab strain *972 h^-^* containing *leu1-32* was used for the transformation of strains, for experiments using *aal1-pOE* and empty-vector control. Two independent deletion strains *aal1Δ1* and *aal1Δ2* were generated by CRISPR-Cas9 using distinct sgRNAs^113^ in the *972 h^-^* strain. Strains *aal1Δ::natMX6* (*aal1Δ*, used for SGA), *kanMX6:P41nmt1-aal1* (*aal1-gOE*, used for smFISH), *rpl1901Δ::kanMX6* (*rpl1901Δ*) and *natMX6:P41nmt1-rpl1901* (*nmt1::rpl1901*, used in Figure 4E and F) were constructed as described^114^ in the *972 h^-^* background and deletion/overexpression primers were designed using the Pombe PCR Primer Programs at http://www.bahlerlab.info/resources/. For the *rpl1901* deletion mutant, the PCR reaction to create the insertion cassettes included 8% DMSO (Invitrogen). The *aal1-pOE* (*P41nmt1-aal1*) and *SPNCRNA.401-pOE* (*P41nmt1-SPNCRNA.401-pOE*) ectopic overexpression strains, driven by the medium-strength *nmt1* promoter^114^, were generated by PCR amplification of the predicted full-length *SPNCRNA.1530* and *SPNCRNA.401* sequences, respectively, as annotated in PomBase^115^, using the high-fidelity Phusion DNA polymerase (NEB) and cloning into the pJR1-41XL vector^116^ with the CloneEZ PCR Cloning Kit (GenScript). Plasmids were checked by PCR for correct insert size and also by Sanger sequencing (Eurofins Genomics) using a universal forward primer (5′ CGGATAATGGACCTGTTAATCG 3′) for the pJR1-41XL plasmid upstream of the cloning site. Plasmids were transformed into *S. pombe* cells (*leu1-32 h-*), and leucine prototroph transformants were selected on solid Edinburgh Minimal Medium (EMM2, with 5g/l NH_4_Cl as nitrogen source). An empty-vector control (evc) strain was created analogously. For *aal1Δ* rescue experiments, the *aal1Δ1* strain was recreated in a *leu1-32 h-* strain and either the *aal1-pOE* or evc plasmid was transformed. All strains were revived in rich YES medium (yeast extract with supplements and 3% glucose) from glycerol stocks. All experiments that included strains expressing genes under the transcriptional modulation of the *P41nmt1* promoter were grown in EMM2 or EMM supplemented with 3.75g/l glutamate (EMMG). All cultures were grown at 32°C with 170 rpm shaking in an INFORS HT Ecotron incubator unless indicated otherwise.

### Quantitative yeast growth assays in liquid culture

The assays were performed in a BioLector microbioreactor (m2p-labs) at 32°C with 1000 rpm shaking and 85% humidity in 48-well microtiter plates (FlowerPlates). The starter cultures were grown to mid exponential phase, diluted in EMMG to achieve initial OD_600_ of ∼0.05 and 1400 µl cultures were incubated in hexaplicates per treatment/genotype with well placements completely randomised to minimise any positional and border effects. For repression of the *P41nmt1* promoter 15 µM thiamine (Sigma) was used. Rapamycin (Cayman Chemical) alone experiments were performed with 300 ng/ml added after 8 hours of initial growth, whereas combined drug experiments used 100 ng/ml rapamycin and 10 mM caffeine (Sigma) added at the start of incubation. Growth (biomass accumulation) was monitored in real time, making measurements every 10 min until the cultures reached stationary phase. The growth data were normalised to time 0. Mean growth curves were fitted and growth rates and lag periods were calculated with *grofit*^117^. For the *aal1-pOE* vs evc +/- thiamine assays, statistical significance was determined with a one-way ANOVA followed by Tukey’s Honest Significant Difference test for all pairwise comparisons, using *R*^118^(ver 4.0.3) (45). Statistical significance of *aal1Δ* against wild-type comparisons were determined with a one-way ANOVA followed by Dunnett’s test (*multcomp*) ^119^.

### Chronological lifespan assays

To determine the CLS, strains driven by a *P41nmt1* promoter and respective controls were grown with or without 15 µM thiamine in EMMG medium whereas deletion mutants and controls were grown in YES medium, each strain as three independent biological replicates. Day 0 was defined as the day the cultures reached a stable maximal cell density. The percentages of viable cells were measured using a robotics-based colony-forming units (CFU) assay^5^. A Poisson distribution-based model was used to obtain the maximum likelihood estimates for the number of CFUs, and percentage viability was calculated relative to that of the CFUs at Day 0 (100% cell survival). CFU measurements were made daily until cultures reached 0.1–1% of the initial cell survival.

### Polysome fractionation and RT-qPCR

Cells were grown in EMMG at 32 °C and 100 ml cultures were sampled for each timepoint. To block translation and capture translating ribosomes, cycloheximide (Sigma) was added to a final concentration of 100 µg/ml and incubated for 5 min with shaking. Cells were collected by centrifugation and lysed in lysis buffer (20 mM Tris-HCl pH 7.5, 50 mM KCl, 10 mM MgCl_2_) supplemented with 100 µM cycloheximide, 1 mM DTT (Sigma), 20 U/ml SuperaseIn (Invitrogen) and protease inhibitors (Complete, EDTA-free, Roche). Lysis was performed with 0.5 mm acid washed beads in a FastPrep instrument (MP, FastPrep24, Settings: speed, 6.0 m/sec; adapter, Quick Prep; time 20 sec; 5 cycles with ≥5 min incubations on ice in between). Number of lysis cycles were increased to ∼12 for stationary phase and aging cells to achieve >80% lysis. The lysates were centrifuged at 17,000 g for 5 min followed by another 15 min at 4°C to remove cell debris, and the lysates were quantified in a Nanodrop (OD_260_). Equal amounts of each lysate were loaded for the polysome fractionation. Then, 10-50% linear sucrose gradients were prepared with a Gradient Master (Biocomp) using 10% and 50% sucrose (Sigma) solutions prepared in lysis buffer freshly supplemented with 100 µM cycloheximide and 1 mM DTT. The lysates were carefully laid on top of the gradients and centrifuged in a SW-41Ti rotor in a Beckman L-80 ultracentrifuge at 35K for 2 h 40 min at 4 °C. The tubes were processed in a Gradient Fractionator (Teldyne ISCO) with 55% sucrose as the chase solution. The polysome fractionation profiles were recorded and the fractions collected (∼800-900 µl per fraction; 12-13 fractions per gradient) and immediately placed on ice. The area under the curve (AUC) of the monosome peak and the polysome peaks were measured with ImageJ^120^ for the calculation of the polysome:monosome ratios (P/M) and total ribosome content (P+M). The P/M was calculated by dividing the sum of the AUC of all polysomes divided by the AUC of the monosome, whereas P+M was the sum of both of these. RNA was extracted with TRIzol reagent (Invitrogen) as per manufacturer’s recommendations. RNA was precipitated with isopropanol overnight at −20 °C. DNase digestion and subsequent RT-qPCR analysis were performed with 1 µg RNA as described below. The calculation of *aal1* and *ppb1* enrichment across polysome profiles was calculated as described^59^. Briefly, the threshold cycle (C_T_) of each fraction was subtracted from the C_T_ of maximum value (always either fraction 1 or 2) for each primer set. The resulting difference in threshold cycles (ΔC_T_) was used to calculate the relative change in mRNA levels between fractions by calculating the 2^ΔCT^ value. The mRNA distribution across the entire polysome profile was graphically presented as the percentage of mRNA in each fraction divided by the total amount of mRNA (sum of 12 fractions).

### RT-qPCR Analysis

RNA was extracted using the TRIzol reagent (Invitrogen) as per manufacturer’s recommendations. In-tube DNaseI (Turbo DNase, Invitrogen) digestion and subsequent reverse transcription (RT) was performed with 1 µg RNA. Random primed cDNA was prepared with SuperScript III reverse transcriptase (Invitrogen) as per standard protocols and all samples had an equivalent RT- reaction. Strand-specific RT with gene specific primers were used for *aal1* transcript quantification with 120 ng/μl Actinomycin D added additionally in the RT reaction. RT-qPCR was performed in a QuantStudio 6 Flex Real-Time PCR System (Applied Biosystems) with Fast SYBR Green Master mix (Applied Biosystems), 1/5 diluted cDNA template and 250 nM primers as per manufacturer’s recommendations. Samples were run in triplicates along with non-template and RT- controls and relative starting quantity was estimated using the ΔΔCt^121^ method. The *aal1* transcript levels across samples were normalized to the *ppb1* expression levels; *ppb1* is the least variable gene under many perturbations including stationary-phase and is comparatively lowly expressed^60^. All other protein-coding transcripts were normalised to *act1* expression levels. Melt curve analysis was performed following amplification to confirm the specificity of amplicons over primer dimers. All primer pairs were initially assessed in a standard curve for efficiencies, and primer pairs with efficiencies of 90-110% were used for RT-qPCR. All primers used are listed in Supplemental Table 1.

### Genome-wide genetic interaction analyses (SGA)

The SGAs were performed as described^48^ using the *S. pombe* Bioneer haploid deletion library v5.0 consisting of 3420 deletion mutants. The *aal1*Δ (*aal1::NatMX6 h^-^*) query strain with a nourseothricin resistance cassette (*Nat*) was generated as described^114^. The *ade6Δ* strain (*ade6::NatMX6 h^-^*), which does not alter the fitness of the mutant library under optimal conditions, was used as a control query strain serving as the equivalent of the fitness of the single mutants^6^. We assessed all pairwise gene interactions using colony size (growth rate) as a measure of double mutant fitness relative to that of the *ade6Δ* control double mutants. The deletion library was revived in YES-agar, grown for 3 days at 32°C and copied on to YES-agar with G418 (100 µg/ml). All query strains including the control were prepared in 384-well format in YES-agar with nourseothricin (100 µg/ml). The query strains were mated with the library mutant strains to create double mutants in 384-well format with a RoToR HDA pinning robot (Singer Instruments) in EMM-N-agar supplemented with adenine, uracil and leucine (100 mg/ml each). The plates were incubated at 25°C for 3 days to allow mating and sporulation followed by a 42°C incubation for 3 days to kill the parental cells. Then, the colonies were copied onto YES-agar and incubated for 1 day to allow spores to germinate. When the colonies were sufficiently grown, they were pinned onto double selection plates: YES-agar with 100 µg/ml G418 and nourseothricin and incubated at 32°C for 2 days. Then, the plates were imaged using a EPSON V800 scanner as .jpg files.

Colony size data were extracted from the images using the R-package *gitter*^122^ (used code: *gitter.batch (image.files=filePath, ref.image.file=reference, plate.format=384, verbose=’l’, inverse=T*). Subsequent analyses were performed in R (ver 4.0.3)^118^. Based on the plate number and row-column position, gene names were assigned to the colony data. Small (< 100 pixels) and absent colonies (extreme outliers) were removed in the *ade6Δ* control plates to avoid false positives arising from absent mutants in the mutant library or differences in pinning and other non-genetic variabilities. These mutants were marked and excluded from subsequent analysis. Then the colony sizes were normalized for spatial effects due to colony position in the plate and plate-to-plate variation by median smoothing and row/column median normalization^123^. Any genes within 250 kb of *aal1* or *ade6* genes were excluded from the subsequent analysis as linked loci. For the rest of the genes with mutants in the library, genetic interaction scores (GIS) were calculated as the log_10_ transformed ratio between the median normalized colony sizes of the experimental query and that of the control. The upper and lower limits of the GIS were arbitrarily set to +2 and −2 for practical reasons and a cutoff of 0.1 was used to call hits.

### Single-molecule RNA fluorescence *in situ* hybridization

smRNA-FISH was performed with antisense probes as described^52^. Since the *aal1* RNA is not detectable during exponential growth and performing smRNA-FISH is technically challenging in aging stationary phase cells^53^, we used the strain overexpressing *aal1* in its native locus (*aal1-gOE*). Cells were grown in EMMG to mid-exponential phase and were fixed in 4% formaldehyde. The cell wall was partially digested using zymolyase. Cells were permeabilized in 70% ethanol, pre-blocked with bovine serum albumin and salmon sperm DNA, and incubated overnight with custom Stellaris oligonucleotide probes (Biosearch Technologies) labeled with CAL Fluor Red 610. Cells were mounted in ProLong Gold antifade mount with DAPI (Molecular Probes), and imaged on a Leica TCS Sp8 confocal microscope, using a 63x/1.40 oil objective. Optical z sections were acquired (0.3 microns z-step size) for each scan to cover the entire depth of cells. The technical error in FISH-quant detection was estimated at 6%–7% by quantifying the *rpb1* mRNA foci with two sets of probes labeled with Quasar 670.

### RNA sequencing

Cells were grown in triplicate per genotype and aged in EMMG and sampled at day 6 of CLS. RNA was extracted with the Qiagen RNeasy Mini Kit. Cells were lysed with 0.5 mm acid washed beads (Sigma) in in a FastPrep instrument (MP, FastPrep24-Settings: speed, 6.0 m/sec; adapter, Quick Prep; time, 20 sec; ∼12 cycles with ≥5 min incubations on ice in between). The amount of beads and the volume of buffer needs to be increased by ∼15% and the number of lysis cycles up to ∼12 for stationary phase, aging cells to achieve lysis of >80% of cells. The cells and beads were loosened by flicking the tubes after every 3-4 cycles to increase the efficiency of lysis. RNA was extracted with RNeasy spin columns and digested with DNase I (Qiagen) for 30 minutes at room temperature followed by a column cleanup with RNeasy spin columns. The rRNA was removed using the Ribo-Zero rRNA removal kit (Illumina) as per recommendations. RNA quality was assessed in an Agilent 2100 Bioanalyzer and strand-specific cDNA libraries were made with NEXTflex™ Rapid Directional qRNA-Seq™ Kit (PerkinElmer, formerly BiooScientific) with molecular indexing. Libraries were quantified in a Qubit, pooled and 4 nM of the pooled library was sequenced in an Illumina HiSeq2500 sequencer, 50 bp paired-end reads in two lanes. All quality control steps including quality filtering, demultiplexing, and adapter removal were performed with Illumina BaseSpace in-house tools leaving unique molecular indices (UMIs) intact. The quality of the sequences was confirmed with FastQC (ver 0.11.2, https://www.bioinformatics.babraham.ac.uk/projects/fastqc/). Reads for each sample from the two lanes were concatenated and the UMIs were clipped and added to the read header by je^124^ (ver. 1.2, http://gbcs.embl.de/je) with the following code: *je clip F1=pair1.fastq.gz F2=pair2.fastq.gz LEN=8*. Reads were aligned with STAR^125^ (ver. 2.6.1a, https://github.com/alexdobin/STAR) against the *S. pombe* genome sequence with the following code: *STAR --genomeDir dir --runThreadN 8 --readFilesIn pair1.fastq.gz pair2.fastq.gz --readFilesCommand gunzip -c --outFileNamePrefix prefix --outSAMtype BAM SortedByCoordinate --quantMode GeneCounts --outWigType wiggle --clip5pNbases 9 -- limitBAMsortRAM 2552288998*. Sorted bam files were used to count the number of reads per gene using htseq^126^ (version 0.12.3, https://htseq.readthedocs.io/en/master/index.html; the code used was: *htseq-count -s reverse -f bam -t gene -i ID file.bam Spombe.gff3 > gene.counts*). The genome sequence and gene annotation gff3 files were downloaded from PomBase^115^. We performed the differential expression analysis with edgeR^127^ (version 3.46.0). Very lowly expressed genes were removed by retaining genes with a mean read count of 1 per million in at least 3 samples. TMM normalization was performed on the filtered counts. Common (overall variability across the genome) and tagwise (measure of the degree of inter-library variation of each gene) negative binomial dispersions were estimated by weighted likelihood empirical Bayes using limma^128^ and fitted in to a generalised linear model with strains as a categorical variable with evc (empty-vector control) strain as the reference. Benjamini and Hochberg method was used to control FDRs (False Discovery Rates).

### ChIRP-MS

The standard ChIRP-MS protocol^55^ was optimized for *S. pombe* cells as described below. Antisense tiling oligo probes for selective pull-down of *aal1* RNA (783 nt) were designed with the online probe designer at *singlemoleculefish.com*, using the following parameters: number of probes = 1 probe/∼100 bp of RNA length; target GC percentage = ∼45%; oligonucleotide length = 20 nt; spacing length = ∼60-80 bp; extensively repeated regions omitted. The probes were checked for homology with other transcripts in the *S. pombe* genome using PomBase Ensembl Blast [options: DNA| DNA database | Genomic sequence | BLASTN | short sequences] and the probes with homology >13bp to any other region (especially cDNA/RNA) were discarded resulting in 5 usable oligo probes out of 8 designed (Supplemental Figure 7). These antisense DNA probes were synthesized with Biotin-TAGs at the 5’ ends (Sigma) and 100 µM probes were pooled in equimolar ratios. The *aal1-pOE*, evc, *aal1Δ* and wild-type cells were grown in 1-litre cultures each in EMMG until Day 6 after entering stationary phase, monitoring the CLS in 1 ml aliquots sampled every other day. On the Day 6, cells were fixed with 37% formaldehyde (3% final, Sigma) for 30 min at room temperature (RT) with gentle shaking (100 rpm), and the reaction was quenched with 2.5 M glycine (0.125 M final, Sigma) for 10 min at RT. Cells were washed once in ice-cold PBS with PMSF (Sigma, 1 mM final, freshly added before use) and the cell pellets were snap-frozen in liquid-N to store at -80° C.

Cell pellets were thawed on ice, and resuspended in ice-cold lysis buffer (50mM Hepes pH 7.6, 1mM EDTA pH 8.0, 150mM NaCl, 1% Triton X-100, 0.1% Na-Doc) with freshly added EDTA-free protease inhibitors (Roche), 1mM PMSF and 100 U/ml SuperaseIn (Invitrogen) in 15 ml tubes. Cells were lysed with 0.5 mm acid-washed glass beads in a FastPrep (MP Biomedicals, FastPrep24-Settings: 15 ml tube adaptors; speed, – 6.0 m/sec; time – 20 sec) for 12 cycles with 5 min incubations on ice in between. Supernatants were collected by centrifugation. Cell lysates were transferred to Diagenode 15 ml sonication tubes and sonicated (Bioruptor pico, Diagenode) 30 sec ON/ 45 sec OFF at 4°C for 60 cycles (6x10), with vortexing the samples after each 10 cycles. Lysates from the same samples were pooled and centrifuged at 16,000g for 10 min at 4°C to collect the supernatants. Then, the lysates were pre-cleared by incubating with Dynabeads (Invitrogen, magnetic Streptavidin) at 37 °C for 30 min with shaking. Before hybridization, beads were removed twice from lysates. For hybridization, 100 pmol of probes in 2 ml hybridization buffer (750 mM NaCl, 1% SDS, 50 mM Tris-Cl pH 7.0, 1 mM EDTA, 15% formamide), supplemented with protease inhibitors, 1mM PMSF and SuperaseIn and incubated at 37°C with shaking overnight. Then, 100 μl Dynabeads per 1 ml lysate were added and incubated at 37°C for 30 min with shaking. The beads were washed with 5 x 1 ml prewarmed (37°C) wash buffer (20 mM Tris-HCl pH 8.0, 140 mM KCl, 1.8 mM MgCl_2_, 10% Glycerol, 0.01% NP-40 with freshly added 1mM PMSF). During each wash, the beads were incubated at 37°C with shaking for 5 min. For protein elution, beads were collected on a magnetic stand, resuspended in biotin elution buffer (12.5 mM biotin – Invitrogen, 7.5 mM HEPES pH 7.5, 75 mM NaCl, 1.5 mM EDTA, 0.15% SDS, 0.075% sarkosyl, 0.02% Na-Doc) and incubated at RT for 20 min and then at 65°C for 10 min with rotation. Eluent were transferred to a fresh tube, and beads were eluted again. The two eluents were pooled, and the residual beads were removed again using the magnetic stand.

TCA was added to 25% of the total volume to precipitate proteins at 4°C overnight. Then, proteins were pelleted at 16,000 g at 4°C for 30 min. The supernatant was carefully removed, the pellets were washed once with cold acetone and air-dried for 1 minute. Proteins were immediately solubilized in 8 M urea and tryptic (Promega) digestion was performed overnight at 37°C with shaking followed by desalting. Peptides were reconstituted in a mixture of 97:3 H_2_O:acetonitrile (containing 0.1% formic acid). The mobile phase consisted of two components: A) H_2_O + 0.1% formic acid and B) Acetonitrile + 0.1% formic acid. Online desalting of the samples was performed using a reversed-phase C18 trapping column (180 µm internal diameter, 20 mm length, 5 µm particle size; Waters). The peptides were then separated using a linear gradient (0.3 µL/min, 35°C column temperature), where Buffer A was transitioned from 97% to 60% over 60 min, on an Acquity UPLC M-Class Peptide BEH C18 column (130 Å pore size, 75 µm internal diameter, 250 mm length, 1.7 µm particle size, Waters). The nanoLC system was coupled online with a nanoflow sprayer and connected to a QToF hybrid mass spectrometer (Synapt G2-Si; Waters, UK) to achieve accurate mass measurements using data-independent mode of acquisition (HDMSE)^129^. Each sample was analysed in technical triplicates, ensuring data reproducibility and reliability. LC-MS grade solvents were used consistently throughout the process: LC-MS H_2_O (Pierce), LC-MS Grade Acetonitrile (Pierce) and LC-MS Formic Acid (Sigma). Lockmass calibration was performed using [Glu1]-fibrinopeptide B (GFP, Waters) at a concentration of 100 fmol/µL. The lockmass solution was introduced through an auxiliary pump at a flow rate of 0.5 µL/min to a reference sprayer, which was sampled every 60 sec.

The acquired data was processed using PLGS v3.0.2 (Waters). The data was queried against an *S. pombe* FASTA protein database (UniProt proteome: UP000002485) concatenated with a list of common contaminants obtained from the Global Proteome Machine (ftp://ftp.thegpm.org/fasta/cRAP). The identification parameters included Carbamidomethyl-C as a fixed modification, and Oxidation (M) and Phosphorylation of STY as variable modifications. To account for incomplete digestion, a maximum of two missed cleavages were allowed. Peptide identification required a minimum of 3 corresponding fragment ions, while protein identification necessitated a minimum of 7 fragment ions. The protein false discovery rate was set at 1%. All identified proteins were tabled in SupplementalDataset2. Differential protein enrichment analysis was performed with DEP^130^. As we observed that imputation of missing values with any available method in the package resulted in false positives, we adopted the following strategy. Any protein not identified in at least 2 out of 3 replicates of at least 1 strain was removed resulting in 218 proteins being identified. The remaining missing values were imputed with 1. The data was background corrected and normalized by variance stabilizing transformation. A stringent cut-off of FDR ≤0.005 and log_2_ ≥6.5 in *aal1-pOE* and/or wild type relative to *aal1Δ* was applied to eliminate *aal1*-RNA-independent background interactions.

### ChIRP-seq

ChIRP-seq samples were identical to those described in ChIRP-MS and the washed beads were reconstituted in equal volume of proteinase K RNA buffer (100 mM NaCl, 10 mM TrisCl pH 7.0, 1 mM EDTA, 0.5% SDS) and reverse cross-linked at 70°C for 1 hr with end-to-end shaking followed by another 1 hr incubation at 55°C after freshly adding proteinase K (Ambion, 5% by volume from 20 mg/ml). Then, the samples were boiled for 10 min at 95°C. RNA was extracted with QIAzol and RNeasy Mini kit (Qiagen) according to the manufacturer’s recommendations, including a DNase treatment. Sequencing libraries were prepared with NEXTflex Rapid Directional qRNA-Seq Kit (PerkinElmer), spiked with 20% PhiX (Illumina) to increase the complexity of the libraries to accommodate clustering and sequenced in an Illumina MiSeq (75 bp, paired-end) instrument. The read processing was as described for RNA-Seq. Reads were mapped with Hisat2^131^ (version 2.2.1, code used: *hisat2 -x genomeIndex --rna-strandness FR --trim5 8 -1 pair1.fastq.gz -2 pair2.fastq.gz -S out.sam --summary-file summary.txt*). sam files were converted to bam files and sorted/indexed with samtools (ver 1.14)^132^ and sorted bam files were used to count reads per gene with htseq^126^ (ver. 0.12.3, https://htseq.readthedocs.io/en/master/index.html; code use: *htseq-count -s reverse -f bam -t gene -i ID file.bam annotation.gff3 > file.count*). The top *aal1*-bound RNAs were determined with edgeR^127^ (version 3.32.1; Supplemental Data 4) from two replicates each of *aal1Δ*, wild type (wt, 972h-) and *aal1-pOE* strains grown and processed independently. Since there were no significantly enriched RNAs (FDR ≤0.05) in the wild-type relative to *aal1Δ* cells, we chose the top 30 enriched RNAs which were verified in IGV^133^ (ver 2.8.0, https://software.broadinstitute.org/software/igv/). For IGV inspection, strand-specific MiSeq reads were processed further with deepTools^134^ (ver 3.5.1, https://deeptools.readthedocs.io/en/develop/) to get normalized gene coverage in counts per million (cpm) per 50 bp bins (bam to bigWig files) for direct comparison.

### *In silico* prediction of *aal1*-*rpl1901* lncRNA-mRNA interaction

*In silico* predictions of *aal1RNA-rpl1901mRNA* interactions were performed with the ViennaRNA Package^135^ (ver. 2.5.0, https://www.tbi.univie.ac.at/RNA/documentation.html) with RNAcofold^136^. RNAcofold tests the probability of RNA-RNA interactions of the provided RNA pairs by computing minimum free energy structures and a base pairing probability matrix. Potential RNA secondary structures were taken into consideration when the pairwise base pairing probabilities and probable interaction sites were predicted. Similarly, the concentration dependency of homo- and heterodimerizations of RNAs were computed for the given input concentrations of the monomers (in mol/lit) and presented as concentration dependency plots of dimerization.

### Cloning and generation of *UAS-aal1* fly line

The *aal1* sequence was PCR amplified from *S. pombe* including the predicted 3’ polyadenylation signal sequence. Using overlap-extension PCR, *UAS,* and *Drosophila HSP-70* promoter sequences were added at the 5’ end, and an additional 150 bp of SV40 polyadenylation signal sequence was added at the 3’ end. The resulting product was transferred to the *pCaSpeR* plasmid by double digestion with *BamHI* and *Not1*, followed by ligation with T4 DNA ligase, and transferred into competent *Escherichia coli*. The correct clone was confirmed by DNA sequencing at Source Biosciences. The plasmid miniprep of the clone was injected into *white^1118^* embryos and randomly integrated into the fly genome using piggyBac transgenesis^137^, at the Department of Genetics Fly Facility (University of Cambridge, https://www.flyfacility.gen.cam.ac.uk/Services/Microinjectionservice/). The *UAS-aal1* fly line with balancer chromosomes was then provided by the Cambridge fly facility. To confirm whether *aal1* was expressed in the fruit fly, we crossed the *UAS-aal1* homozygous males with *daughterless GAL4* virgins. The flies emerging from this cross were collected and RNA was extracted from five 7-day-old adult flies, using TRIzol (Invitrogen) according to the manufacturer’s protocol. cDNA was synthesized using random hexamers and Superscript II (Invitrogen) according to the manufacturer’s instructions. Using this cDNA as a template, the expression of *aal1* RNA was then confirmed by PCR using primers listed in Supplementary Table 1.

### Fly husbandry and maintenance

We used an outbred wild-type stock that was initially collected from Dahomey (present Benin) in 1970 and subsequently maintained in large population cages on a 12hr:12hr light/dark cycle at 25°C to maintain lifespan and fecundity at levels similar to wild-caught flies. The *white^1118^* mutation was introduced into this background to allow easier tracking of transgenes and Wolbachia infection was cleared by tetracycline treatment. Before the experiments, the *aal1* fly lines and gene-switch drivers (TIGS) were backcrossed into this white Dahomey (wDah) background for at least six generations. All stocks were maintained, and experiments were conducted at 25°C and 60% humidity with 12hr:12 hr light/dark cycles, on SYA food containing 10% brewer’s yeast, 5% sugar, and 1.5 % agar with nipagin and propionic acid added as preservatives.

### Lethality test in flies

To test whether *aal1* exhibits lethality when expressed during development, we crossed *UAS-aal1* homozygous males with daughterless *GAL4* virgins. The adults were allowed to mate for 48 hr, and eggs were collected within 24 hours and counted. After 10 days, the number of flies that emerged from the resultant crosses was recorded. We used GAL4-alone and UAS-alone controls in parallel.

### Lifespan analysis in flies

For lifespan assays, experimental flies were generated from suitable crosses in cages containing grape juice, agar, and live yeast. Flies were allowed to mate and the eggs were collected after 22 hrs and 20 μL of egg sediments (in 1xPBS) were seeded on SYA medium in glass bottles to rear flies at standardized larval densities. Flies emerged after 10 days, and were transferred to new bottles where they were allowed to mate for 48 hrs before sorting females into experimental vials at a density of 15 flies per vial. To induce transgene expression using the GAL4/UAS GeneSwitch system, RU486 (Sigma, dissolved in ethanol) was added to the media at a final concentration of 200 µM. As a control (RU-), equivalent volumes of the vehicle alone were added. Flies were transferred to fresh vials three times a week and their survival scored. To control for potential RU_486_ artifacts, driver-only controls feeding RU_486_ were included in the experiment.

## Data Availability

ChIRP-MS data have been submitted to the ProteomeXchange Consortium via the PRIDE partner repository with the dataset identifier PXD045625. ChIRP-seq and RNA-seq data have been submitted to GEO under the accession number GSE243036.

## Acknowledgments

We thank Melania D’Angiolo and Manuel Lera-Ramirez for their helpful comments on the manuscript. This work was supported by the Biotechnology and Biological Sciences Research Council (Grant numbers BB/R018219/1 to J.B. and BB/W013525/1 to N.A.), the Medical Research Council (to S.M.), and a Bangabandhu Overseas Scholarship, University of Dhaka, to A.F.S.

## Supplemental Files

### Supplemental Figures

**Supplemental Dataset 1:** Data related to SGA assays

**Supplemental Dataset 2:** Data related to ChIRP-MS experiments

**Supplemental Dataset 3:** Data related to RNA-seq of *aal1-pOE* cells

**Supplemental Dataset 4:** Data related to ChIRP-seq experiments

**Supplemental Table 1:** Primers for RT-qPCR, RT-PCR and PCR (*Drosophila*)

**Supplemental Figure 1:**
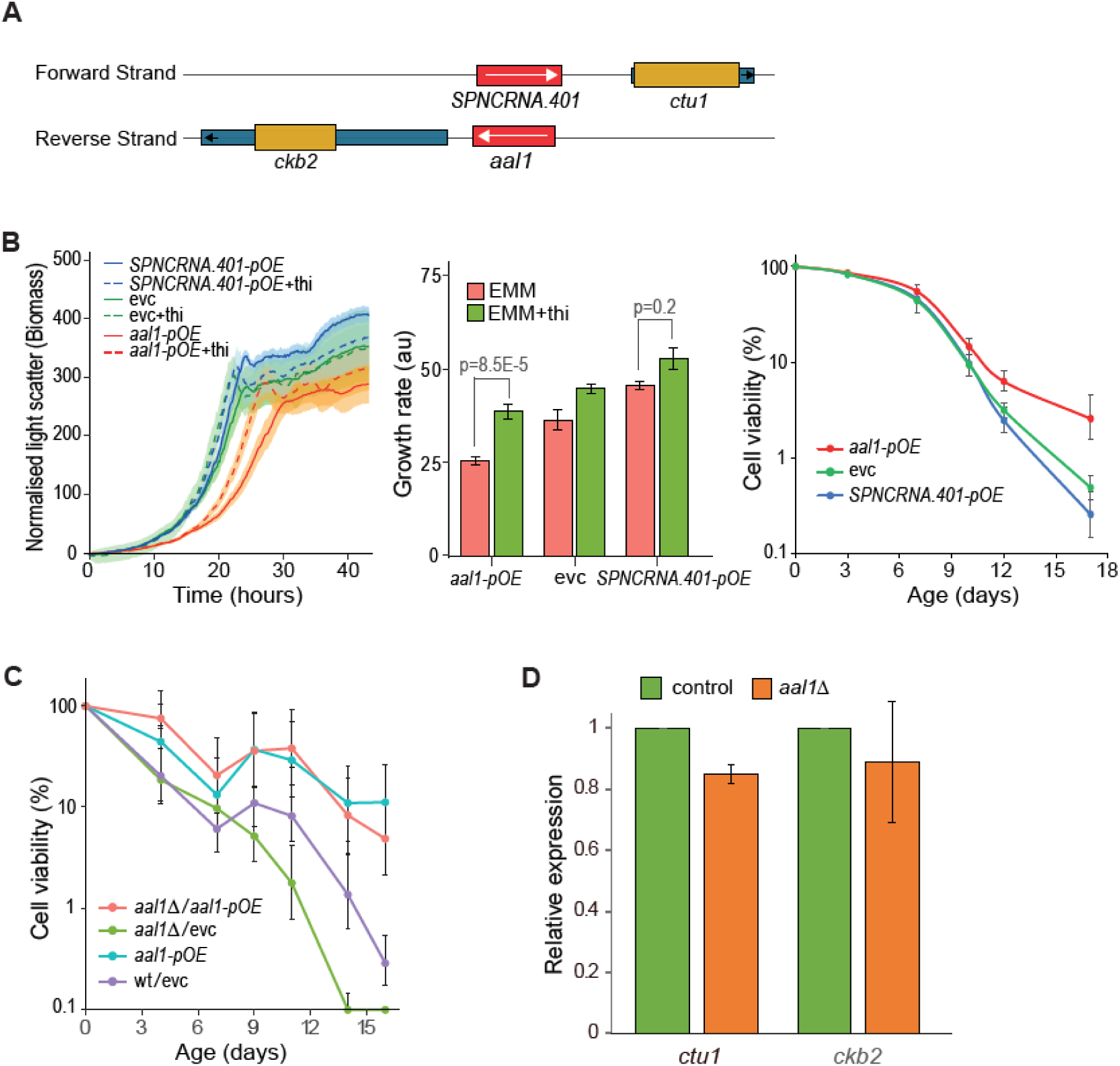
Analysis of the *SPNCRNA.401* lncRNA. **(A)** Genomic environment of *aal1* gene showing a 745 nucleotide overlap in antisense direction with the *SPNCRNA.401* gene. *Red boxes*: lncRNA genes, with transcriptional direction indicated by white arrows; *ochre and blue boxes*: open reading frames and untranslated regions, respectively, of coding genes. Visualization using the PomBase genome browser^139^. **(B)** *Left graph*: Growth assay for cells ectopically overexpressing *aal1* and *SPNCRNA.401* under the thiamine-repressible *P41nmt1* promoter (*aal1-pOE* and *SPNCRNA.401*-*pOE*) compared to empty-vector control (evc) with/or without 15 µM thiamine (thi) added to the medium. Experimental setup and analysis as in Figure 1D. *Middle graph:* Quantitation of growth rate for experiments shown in the left graph, as in Figure 1D. *Right graph:* CLS assays for *aal1-pOE* and *SPNCRNA.401*-*pOE* cells compared to empty-vector control (evc) cells. Experimental setup and analysis as in Figure 1C. **(C)** CLS assays for *aal1Δ* cells ectopically overexpressing *aal1* (*aal1Δ/aal1-pOE*) compared to *aal1Δ* and wild-type cells overexpressing empty-vector controls (*aal1Δ/*evc; wt/evc) and *aal1-pOE* cells. **(D)** Expression of genes flanking *aal1* in the presence and absence of *aal1*. Quantitative RT-PCR experiment to determine transcript levels of *ctu1* and *ckb2* in *aal1Δ* cells relative to wild-type cells (control). Four independent biological repeats were carried out using 7-day old stationary-phase cells. Data were normalized to *act1* expression. The expression of *ctu1* was slightly lower in *aal1Δ* compared to control cells (p_Student’s T_ ∼0.001), while the expression of *ckb2* showed no significant difference (p_Student’s T_ ∼0.35).

**Supplemental Figure 2:**
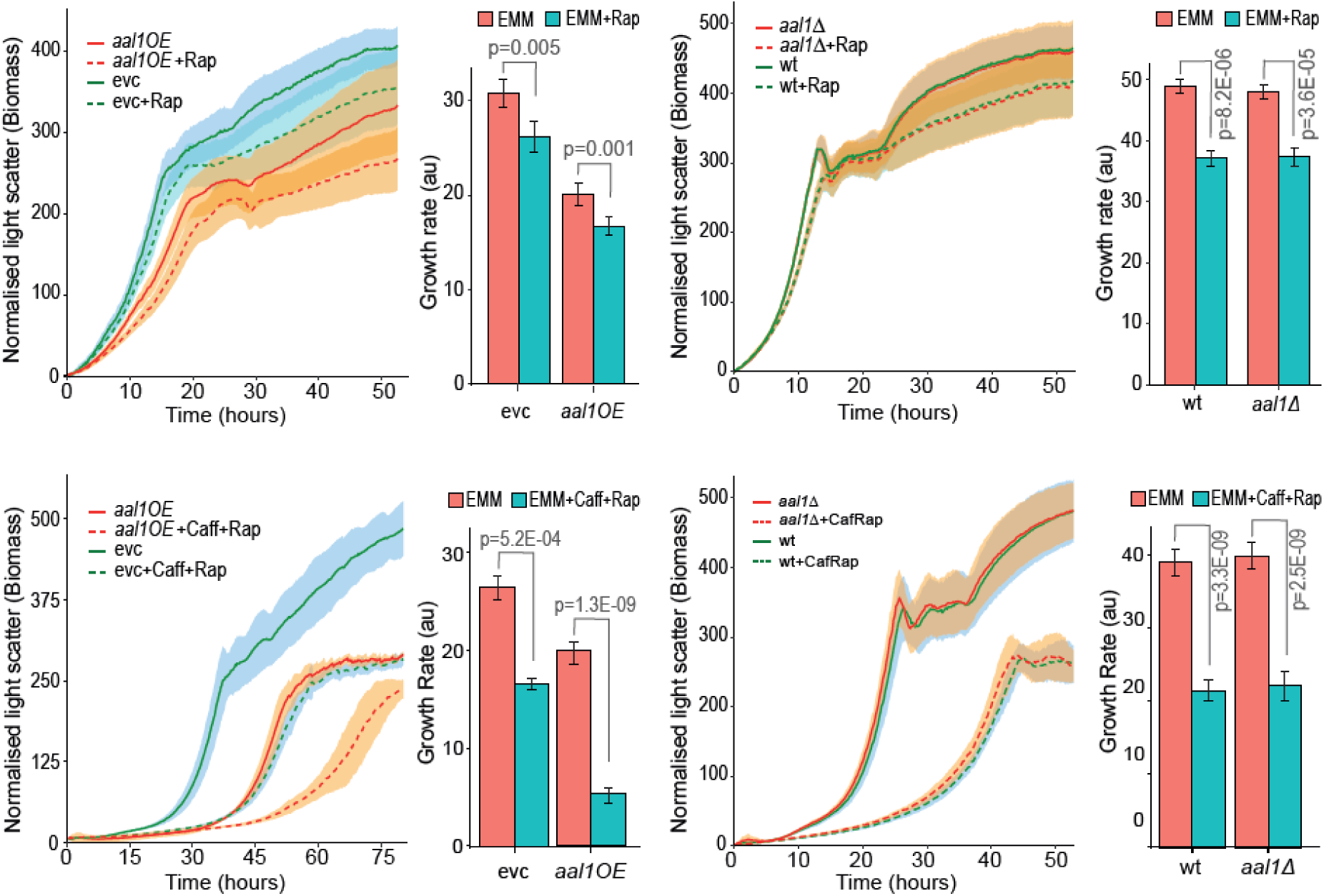
*aal1* phenotypes do not depend on TORC1 signalling. *Top graphs:* Rapamycin inhibits cell growth in *aal1-pOE* and *aal1Δ* mutants to a similar degree as in the respective controls, indicating that TORC1 and *aal1* functions exert additive effects. To avoid overly long lag periods, rapamycin (300 ng/ml) was added after 8 hours of initial growth. *Bottom graphs:* The combination of caffeine (10 mM) and rapamycin (100 ng/ml) leads to a stronger inhibition of cell growth in *aal1-pOE* and *aal1Δ* mutants, similar as in the respective controls, indicating again that TORC1 and *aal1* functions have additive effects. Cells were grown in a microbioreactor and mean growth curves were fitted with *grofit*^115^, with SD shown as shades. Quantitation of growth rates (mean ± SE) for experiments is shown in the bar graphs. Details as in Figure 1D,E.

**Supplemental Figure 3:**
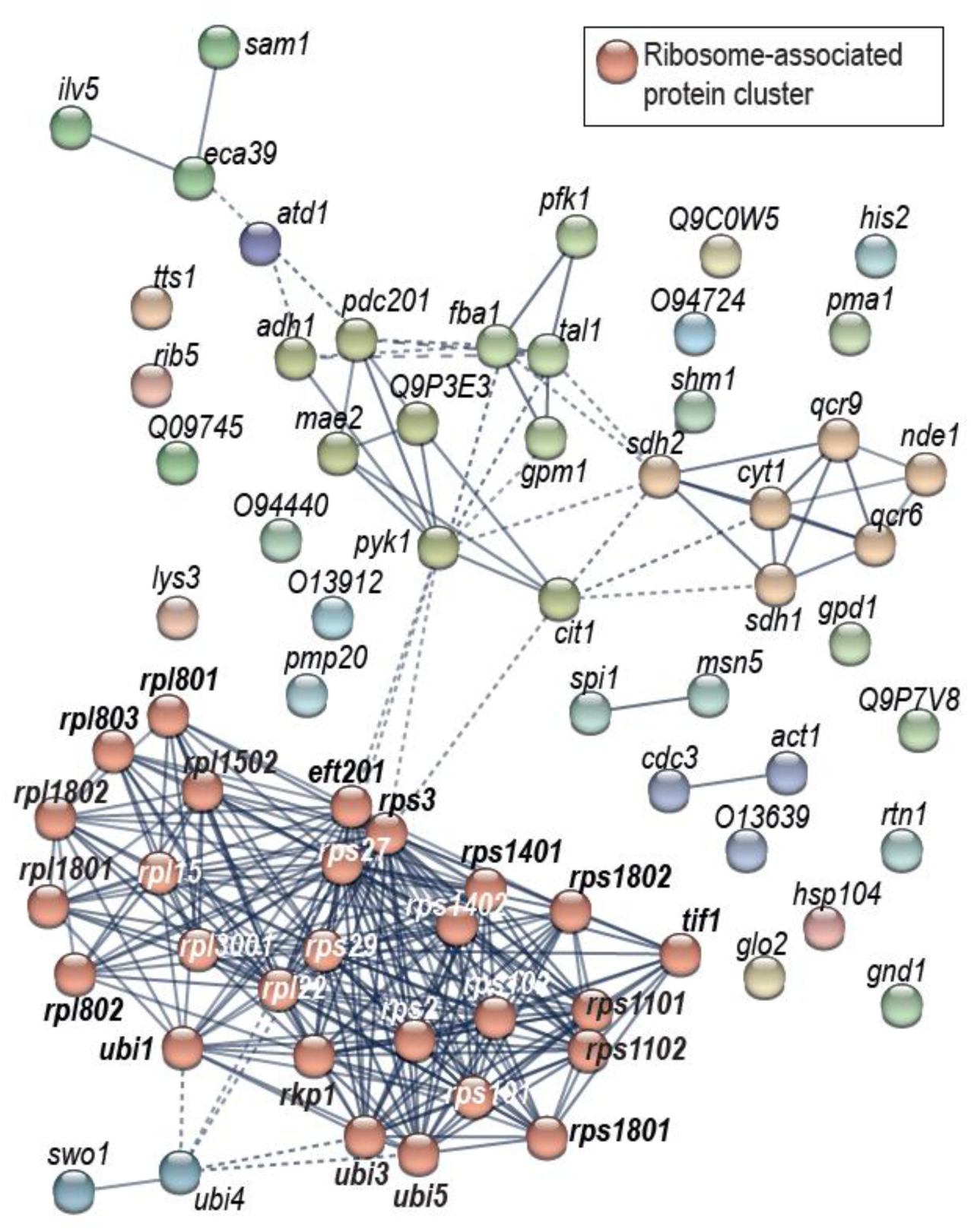
The 68 *aal1-*bound proteins are more highly connected with each other than expected by chance based on known protein-protein interactions. Analysis of protein-protein interaction network with STRING^140^ for the 68 *aal1-*bound proteins reveals more connections than expected by chance (p=1.0e-16). The following STRING parameters were used: active interaction sources = only experiments and databases; minimum required interaction score = high confidence (0.70); cluster with MCL; inflation=3. Dashed lines indicate lower confidence interactions.

**Supplemental Figure 4:**
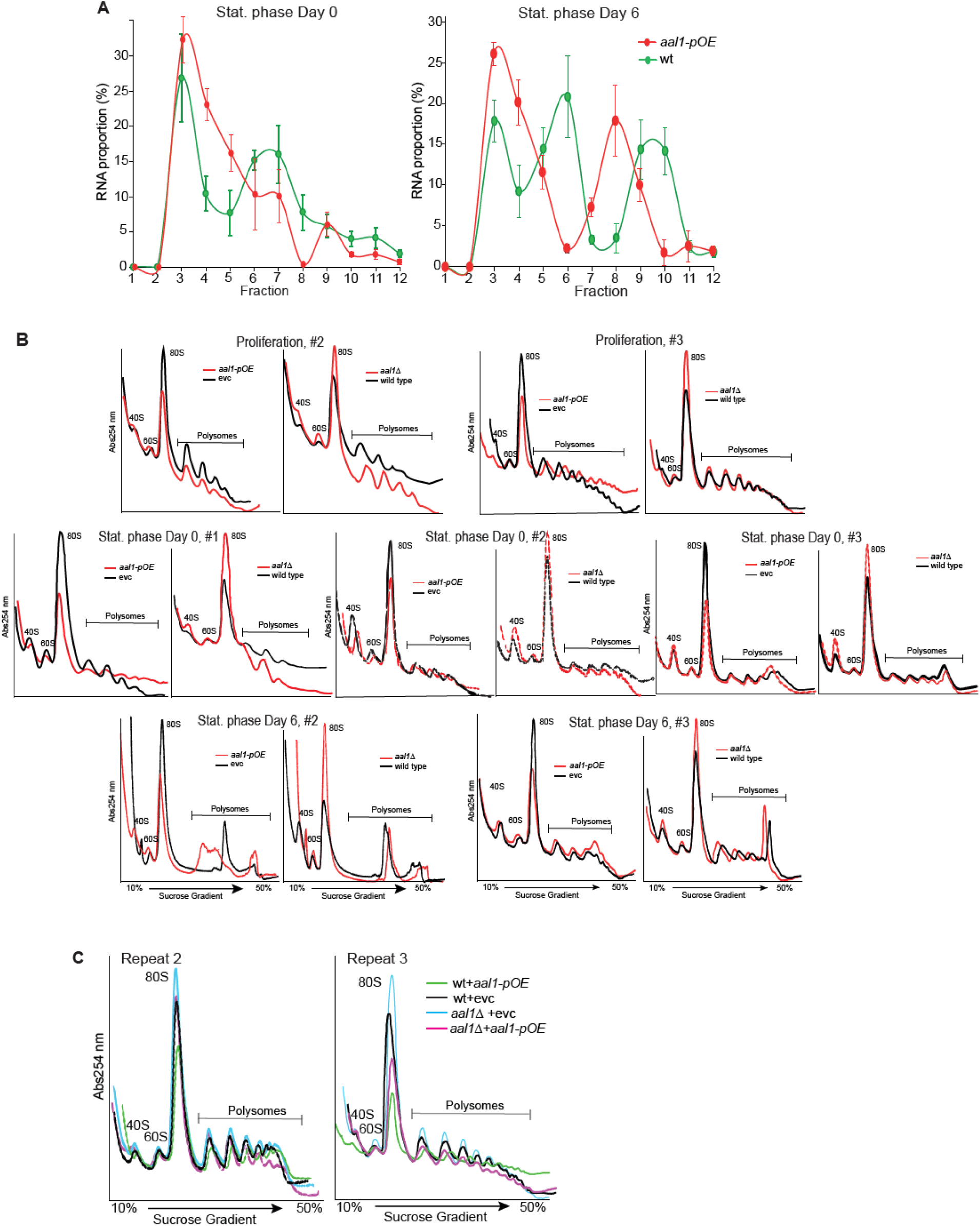
*aal1* associates with ribosomes and reduces the cellular ribosome content. **(A)** Polysome fractionation followed by RT-qPCR shows that *aal1* binds to ribosomes during early stationary phase (Day 0, *left graph*) and late stationary phase (Day 6, *right graph*) in both wild-type (green) and *aal1-pOE* (red) cells. **(B)** Independent biological repeats of polysome profiling as in Figure 3C-F for proliferating cells (top), early stationary-phase cells (middle) and late stationary-phase cells (bottom) for the four strains indicated. The two profiles are aligned at the lowest points of the monosome peaks, corresponding to the baseline. **(C)** Two independent biological repeats of experiment shown in Figure 3G.

**Supplemental Figure 5:**
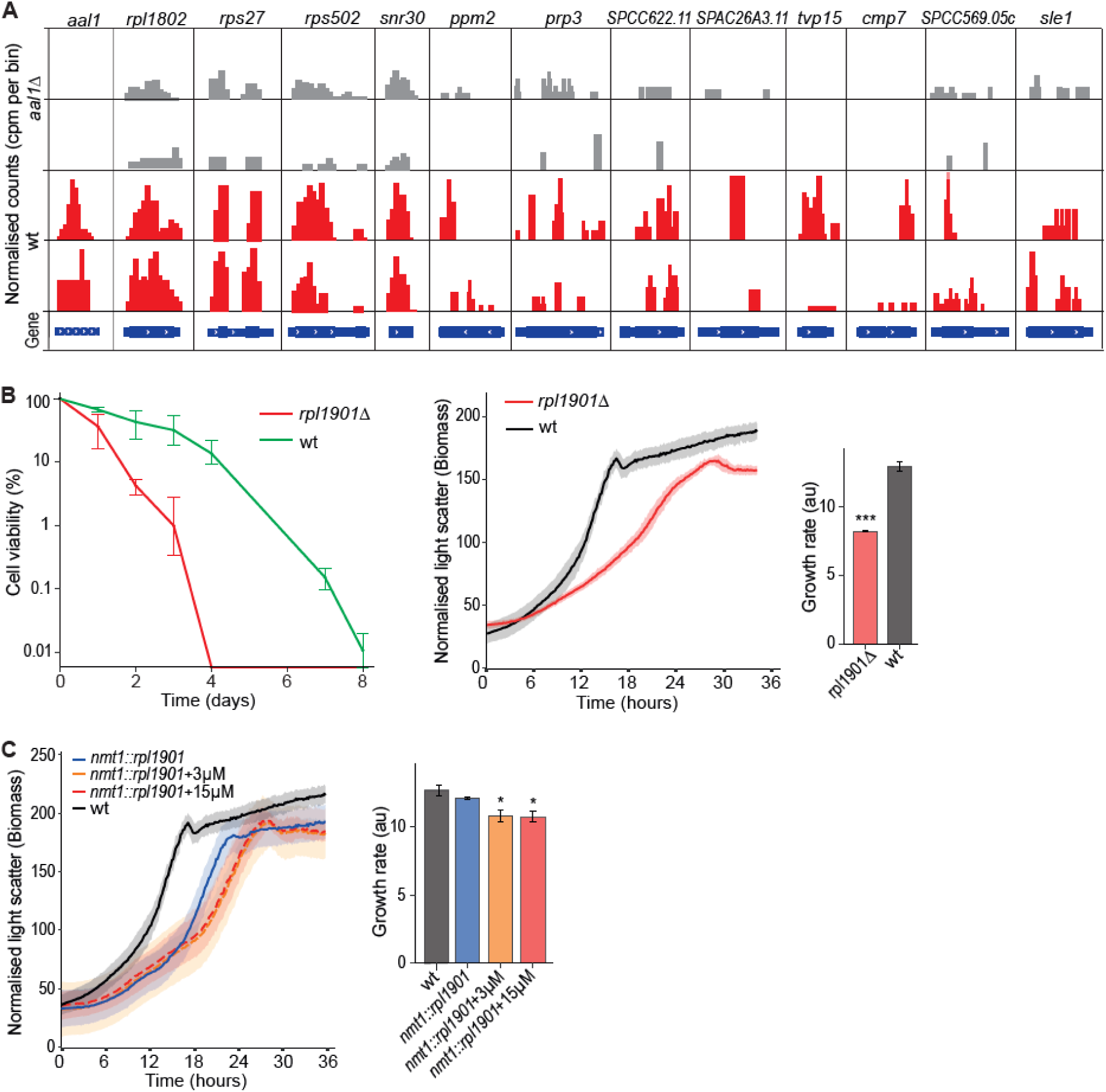
Analyses of RNAs binding to *aal1*. **(A)** IGV tracks from strand-specific ChIRP-seq reads in counts per million (cpm) per 50 bp bins (deepTools)^133^ across *aal1* (control) and prospective target RNAs as indicated on top. Top *aal1*-bound RNAs were determined with edgeR^126^ (Supplemental Dataset 4) from two replicates each of *aal1Δ*, wild type (wt) and *aal1*-*pOE* cells, and the data were verified in IGV^132^ (details in Methods). The prospective *aal1*-bound RNAs in the ChIRP-seq data include three additional mRNAs encoding ribosomal proteins and one small nucleolar RNA (*snr30*). **(B)** *Left graph*: Chronological lifespan assays for *rpl1901Δ* and wild-type cells, performed in rich medium. *Right graphs*: Growth assays of *rpl1901Δ* and wild-type cells and quantitation of growth rate for these assays. Experimental setup and analysis as in Figure 1D. Statistical significance was determined with one-way ANOVA followed by Dunnett’s test^117^, with p <0.0001 relative to wt. **(C)** Growth assays of *nmt1::rpl1901* and wild-type cells with addition of different doses of thiamine as indicated, and quantitation of growth rate for these assays. Experimental setup and analysis as in Figure 1D. Statistical significance was determined with one-way ANOVA followed by Dunnett’s test^117^, with p <0.006 relative to wt.

**Supplemental Figure 6:**
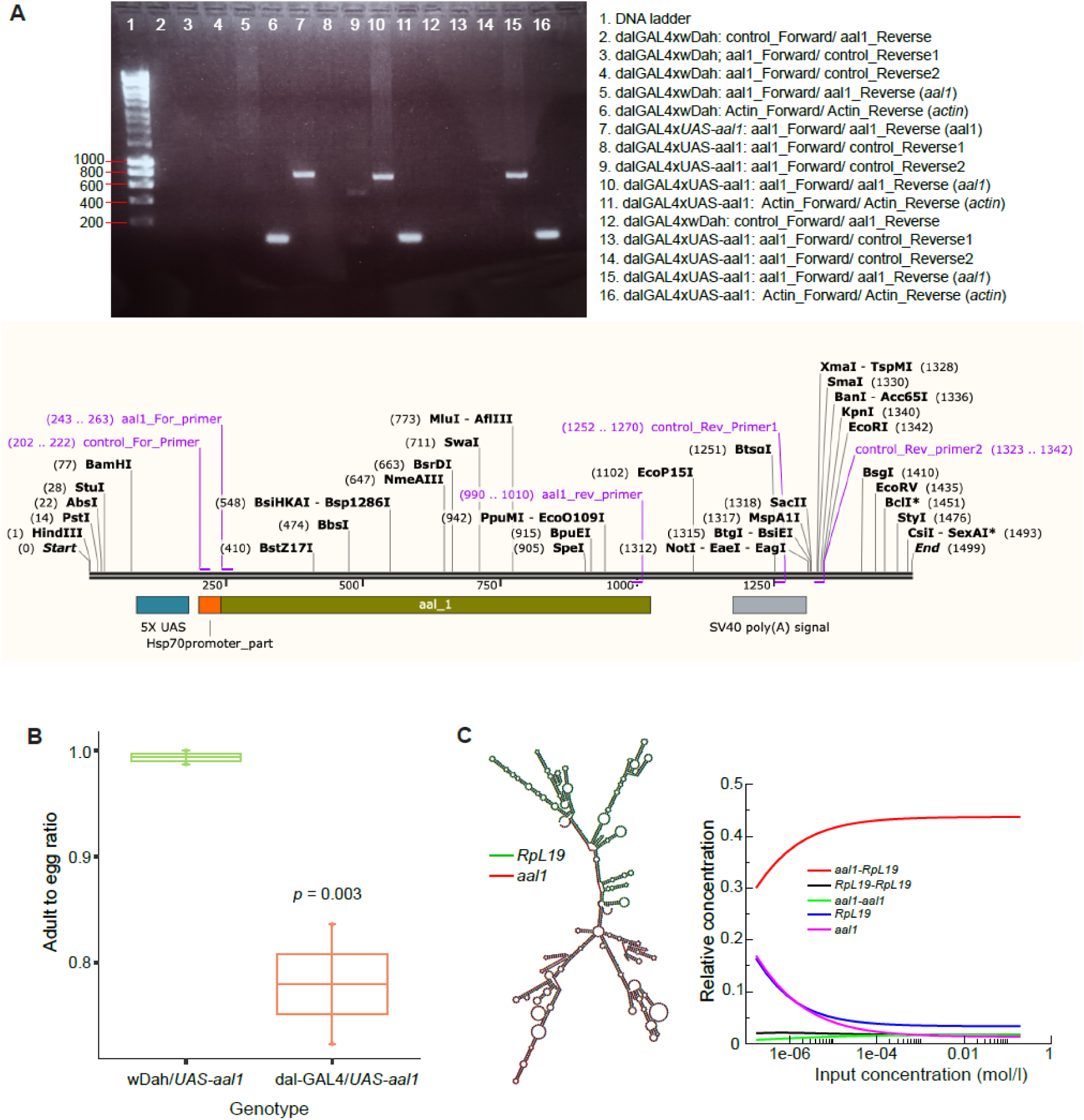
Supporting analyses for experiment expressing *aal1* in flies. **(A)** *Top:* Confirmation of *aal1* expression in the inducible *UAS-aal1* strain flies. The expression of an RNA of expected size (783 nt) was confirmed in females where *UAS-aal1* was driven by the ubiquitous, constitutive GAL4 driver (*daughterlessGAL4 / dalGAL4*). Random primed cDNA was used as template. Primer positions as follows (see scheme). Forward control: resides in Hsp70 promoter; Reverse_control1: in SV40 polyA signal; Reverse_control2: in downstream sequence of SV40 polyA signal; aal1_Forward & aal1_Reverse: in 5’ and 3’ ends of *aal1* transcript, respectively; Actin_Forward/Actin_Reverse: housekeeping Actin gene. wDah control crossed to dalGAL4 was used as a negative control strain (Lanes 2-6 and 11). Three *UAS-aal1* replicates were tested (lanes 7, 8-11, 13-16). Primer sequences are provided in Supplemental Table 1. *Bottom:* Scheme of *UAS-aal1* construct showing the positions of the primers used (purple), visualized with SnapGene Viewer 5.3.2. **(B)** Ubiquitous expression of *aal1* in flies with a *dalGAL4* promoter throughout development results in a significant reduction in the number of flies that reach adulthood (details in Methods: Lethality test in flies). Statistical significance determined with two-sample t-test. **(C)** *In silico* predictions of interation between *S. pombe aal1* and *Drosophila RpL19* using the ViennaRNA package^134^ with RNAcofold^135^. *Left*: Predicted *aal1*-*RpL19* heterodimer structure showing the interaction sites along with potential RNA secondary structures. *Right*: Concentration dependency plot of dimerization showing the computed homo- and hetero-dimerizations of RNAs for concentration relative to each other (y-axis) and different input concentrations (x-axis), with predicted equilibrium concentrations for the two monomers, *aal1* and *RpL19,* the two homodimers, *aal1-aal1* and *RpL19-RpL19,* and the *aal1-RpL19* heterodimer.

**Supplemental Figure 7:**
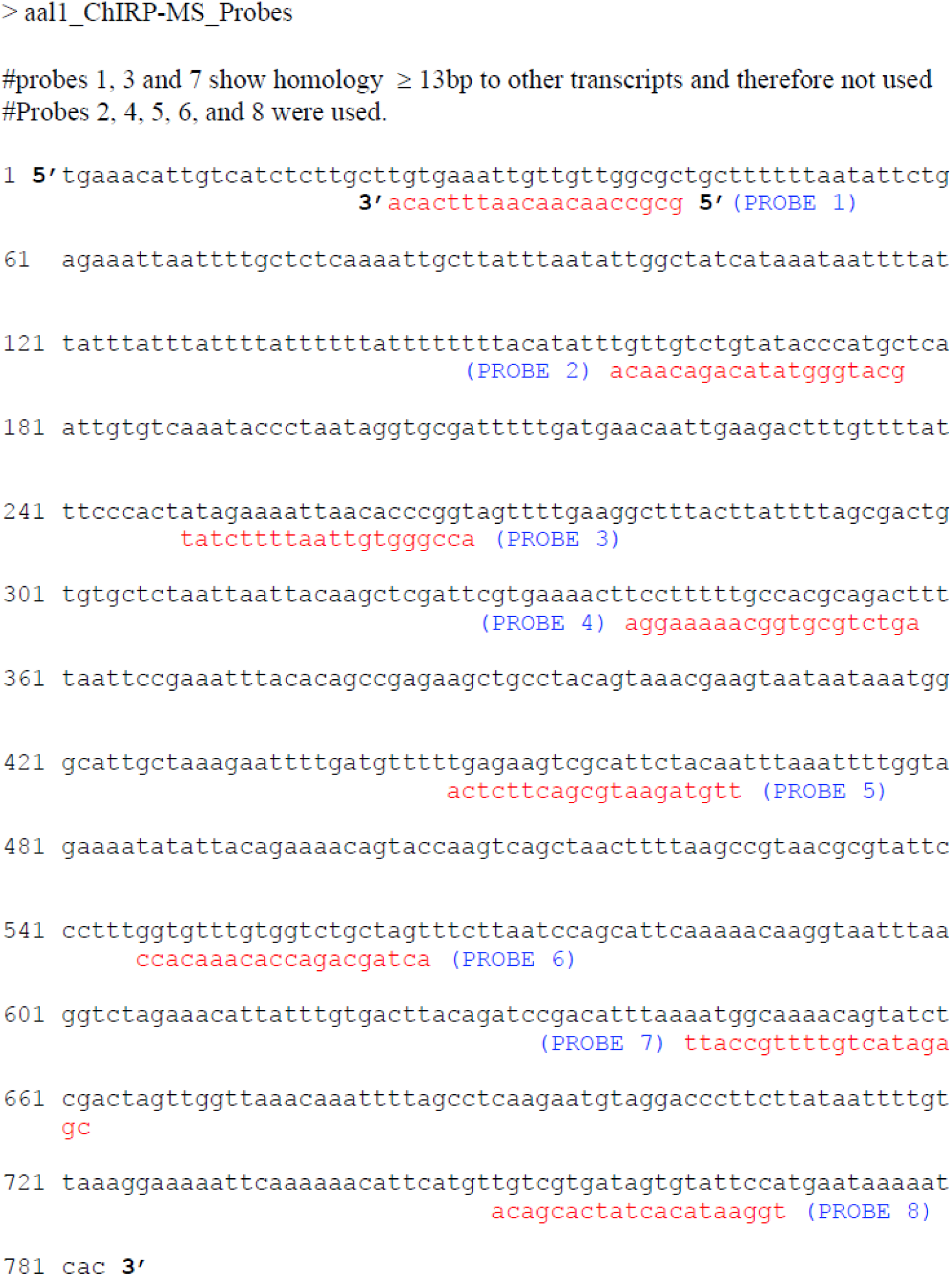
Biotinylated antisense oligo probes used to pull-down *aal1* in ChIRP experiments.

**Supplemental Table 1:**
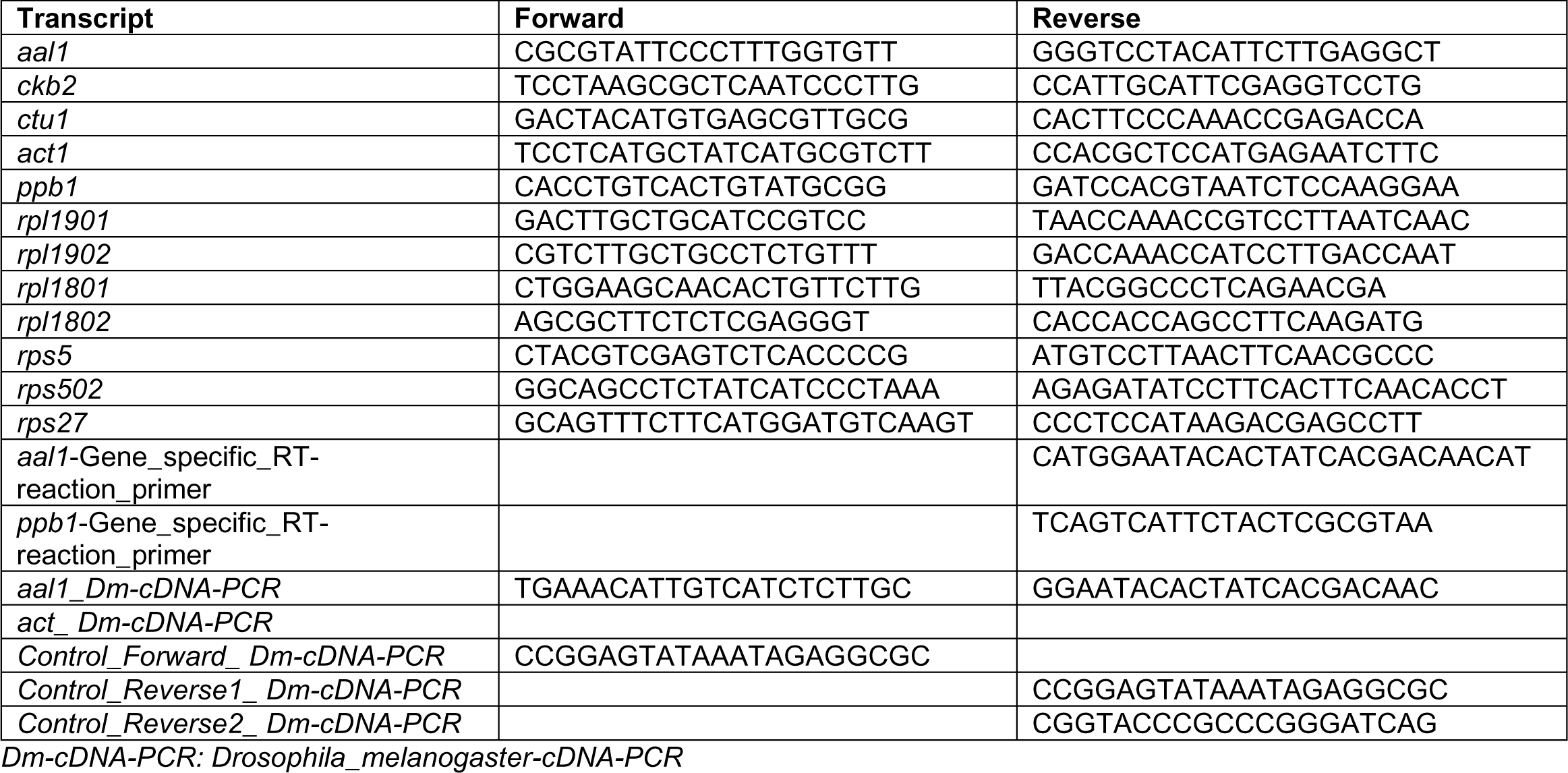
Primers for RT-qPCR, RT-PCR and PCR (*Drosophila*)

